# Replication-stress response shapes *Leishmania* genome plasticity through non-equivalent RAD9 and HUS1 functions

**DOI:** 10.64898/2026.11.08.765153

**Authors:** Ricardo Obonaga, Jennifer A. Black, Stela Virgilio, Richard McCulloch, Jeziel D. Damasceno, Luiz R. O. Tosi

**Affiliations:** Department of Molecular and Cellular Biology, Ribeirão Preto Medical School, University of São Paulo, Ribeirão Preto, 14040-900, SP, Brazil; University of Glasgow Centre for Parasitology, School of Infection and Immunity, Sir Graeme Davies Building, 120 University Place, Glasgow, G12 8TA, United Kingdom; Department of Biochemistry and Immunology, Ribeirão Preto Medical School, University of São Paulo, Ribeirão Preto, 14040-900, SP, Brazil

## Abstract

Genome plasticity, manifested as gene copy-number variation and mosaic aneuploidy, enables adaptation in *Leishmania*, yet how genome-surveillance pathways shape this variation remains unclear. Here we show that partial depletion of HUS1 or RAD9, subunits of the conserved 9-1-1 checkpoint clamp, produces opposing replication-stress responses and contrasting patterns of genome remodelling in *Leishmania major*. During methotrexate selection HUS1 deficiency restricted *DHFR-TS* amplification and favoured linear amplicons, whereas RAD9 deficiency was associated with extensive circular amplification. This divergence extended to genome-wide scale: HUS1-deficient cells carried fewer but larger candidate rearrangements, whereas RAD9-deficient cells retained a largely wild-type rearrangement burden. These differences were accompanied by distinct HUS1 and RAD9 occupancy profiles around repetitive DNA implicated in gene amplification. Deficiency of either of these factors also resulted in differing effects on aneuploidy. While HUS1-deficient cells showed limited shifts towards chromosome disomy, RAD9-deficient cells exhibited broader chromosome copy-number gains and losses. Together, these findings identify replication-stress surveillance as a determinant of parasite genome plasticity, revealing that perturbing individual subunits of the same checkpoint complex can produce opposing genome-remodelling outcomes and reshape the variation available for adaptation.

## Introduction

Faithful genome inheritance requires accurate DNA replication and preservation of genome organization, including sequence integrity, gene and chromosome copy number and whole-genome ploidy. Yet genomes are not invariant: alterations in sequence, architecture and ploidy can be programmed components of normal physiology (1, 2) or selectively retained adaptations (3). Examples include programmed DNA rearrangements that generate immune diversity in vertebrates and genome remodelling or aneuploidy that facilitates adaptation in microbial eukaryotes (3–5). The capacity to generate genomic variation is therefore an important feature of many biological systems, but it must be constrained to preserve genome function and viability.

By contrast, failure to constrain genome variation can produce genome instability, manifested as an increased propensity to acquire mutations, structural rearrangements, copy-number variations (CNVs) and aneuploidy (6, 7). These alterations can reduce cellular fitness and are major contributors to genetic disease and cancer (8). Although their effects are context-dependent, aneuploidies are frequently deleterious because gene-dosage imbalances disrupt the stoichiometry of multiprotein complexes and impose proteotoxic and metabolic stress (8, 9). The balance between potentially beneficial variation and harmful genome instability is therefore largely determined by the pathways that coordinate DNA replication, repair and cell-cycle progression (10).

Accurate DNA replication requires replication forks to traverse the genome efficiently yet forks routinely encounter obstacles that slow or stall their progression, a condition broadly termed replication stress (RS). Major sources include nucleotide depletion or imbalance, DNA lesions, tightly bound DNA-protein complexes, stable DNA secondary structures and conflicts between the transcription and replication machineries (11). Such impediments can uncouple DNA synthesis from helicase progression, exposing stretches of single-stranded DNA (ssDNA) and, if unresolved, can lead to fork collapse and DNA breaks or allow under-replicated regions to persist into mitosis (11). These outcomes can promote chromosome breakage and rearrangement, CNVs and aneuploidy (12, 13). Cells have therefore evolved RS response pathways that detect and stabilize perturbed forks, restrain cell-cycle progression and coordinate fork repair and replication restart. In most eukaryotes, the cellular response to RS is mediated via the atypical kinase ATR, activated in response to RPA-coated ssDNA (11, 14). In parallel, the ring-shaped RAD9-RAD1-HUS1 (9-1-1) clamp is loaded at 5′-recessed ssDNA-dsDNA junctions (15) and contributes to checkpoint signalling and the coordination of downstream repair processes alongside ATR (14, 15). Thus, effective RS surveillance acts to prevent replication-associated lesions from becoming sources of genome instability, whereas altered processing or persistence of these intermediates may contribute to the generation of genome variation.

Against the usual association between extensive genome variation and compromised fitness, *Leishmania* spp. stand out as an unusual example. These unicellular eukaryotic parasites cause leishmaniasis, a spectrum of diseases affecting humans and other mammals, and undergo a digenetic life cycle alternating between phlebotomine sand fly vectors and mammalian hosts (16). More than 20 *Leishmania* species cause human disease in cutaneous, mucocutaneous and visceral forms. Leishmaniasis is endemic across tropical and subtropical regions and remains a substantial global health burden. According to the World Health Organization, more than one billion people live in endemic areas, and an estimated 700,000 to 1 million new cases occur annually (17). Genome plasticity is a hallmark of *Leishmania* biology, manifesting across at least two distinct genomic scales. First, direct and inverted short, interspersed degenerate retroposons (SIDER1 and SIDER2) distributed throughout the genome mediate genomic DNA rearrangements, generating intrachromosomal amplifications as well as circular or linear extrachromosomal amplicons that encompass multiple genes (18). Second, chromosome copy number varies both between chromosomes within individual cells and among cells across a population, rendering mosaic aneuploidy a widespread phenomenon rather than a rare aberration (19–21).

Together, these alterations generate standing genetic variation within parasite populations, enabling the selection of karyotypes or local CNVs that confer a fitness advantage under changing environmental conditions. This is especially relevant in the context of gene expression modulation in this parasite. In *Leishmania*, protein-coding genes are largely intron-less, lack conventional gene-specific RNA polymerase II promoters and are organised into large polycistronic transcription units (PTUs), transcribed as multigene precursors (22). Gene expression is therefore regulated predominantly at the post-transcriptional level with genome plasticity providing an additional means of modulating expression by altering gene dosage (23). In *L. tropica*, gene dosage explained more than 85% of expression variation among genes exhibiting at least twofold differences in transcript abundance (23). Similarly, in *L. donovani*, mean chromosome-wide transcript and protein abundance scaled with somy, although protein-level dosage compensation attenuated this effect for subsets of proteins, notably components of multiprotein complexes (24). Genome plasticity also shapes clinically important phenotypes and frequently accompanies the emergence of drug resistance. Notably, loss of the *L. infantum* miltefosine-sensitivity locus from infecting isolates was associated with a 9.4-fold greater risk of miltefosine treatment failure (25). More broadly, genome-wide analysis of *L. infantum* isolates from patients with visceral leishmaniasis in Brazil identified 17 parasite CNVs significantly associated with patient mortality (26).

Although the phenotypic consequences of *Leishmania* genome plasticity are increasingly well established, the genome-maintenance pathways governing when and where this variation arises remain poorly defined. Homologous recombination and DNA end-processing factors, including RAD51, RAD51-4, and MRE11, influence the formation of locus-specific CNVs, whereas recent studies have linked R-loop homeostasis and ATR-dependent replication timing to chromosome copy number variation and broader genome instability (27, 28). Despite these advances, whether and how the RS response controls the hallmark forms of *Leishmania* genome plasticity remains unclear. In *L. major,* RAD9, RAD1, and HUS1 associate to form the 9-1-1 clamp, but RAD9 is also found in 9-1-1-independent complexes, whereas HUS1 exists as a monomer, suggesting compartmentalised functions for these subunits within this pathway (29). Consistent with this, HUS1 and RAD9 deficiencies produce markedly different effects on DNA synthesis, cell-cycle progression, and recovery from RS (29). Replication timing of subtelomeres, which have been proposed to replicate beyond the time constraints of S-phase, is affected mainly by RAD9 deficiency, whereas only a minor effect is seen upon HUS1 deficiency (30). HUS1 also has a complex relationship with genome stability in *L. major*: its depletion leads to increased SNP accumulation under non-stressed conditions, contrasting with reduced SNP accumulation upon chronic RS, revealing that 9-1-1-associated functions extend beyond the canonical preservation of genome integrity (31). Together, these findings indicate that 9-1-1 components have independent and specialised roles in *Leishmania* genome maintenance, but whether their distinct responses to RS translate into differential control of genome plasticity across different genomic scales remains unresolved.

Here, we investigated how reduced levels of RAD9 or HUS1 reshape both local CNVs and mosaic aneuploidy in *L. major*. We show that HUS1 and RAD9 deficiencies produce distinct patterns of genome variation: HUS1 deficiency primarily restricted the accumulation of circular extrachromosomal amplicons under methotrexate selection as well as genome-wide rearrangements, whereas RAD9 deficiency produced broader changes in chromosome somy. These divergent outcomes were accompanied by distinct patterns of ssDNA accumulation and differential RAD9 and HUS1 occupancy at genomic repeats associated with extrachromosomal amplicon formation. Together, our findings establish that RS surveillance contributes to the spectrum and genomic context of genome variation in *Leishmania*, linking the cellular response to RS with the emergence and persistence of diverse forms of genome plasticity.

## Results

### Deficiency of HUS1 and RAD9 produces opposing cell-cycle responses to replication stress

To examine HUS1 and RAD9 roles in genome-plasticity phenotypes that may emerge or be selected over many generations, we required mutant lines that could be propagated long term. We previously showed that HUS1 (31) and RAD9 (29) are essential for *L. major*. *HUS1*-null parasites generated through inducible gene deletion remained viable only briefly (31), precluding their use in long-term assays of CNV and mosaic aneuploidy. Because null mutants of *HUS1* or *RAD9* were not available, we used viable heterozygous lines (*HUS1^+/−^* and *RAD9^+/−^*) allowing us to examine the consequences of reduced HUS1 or RAD9 abundance on genome plasticity over long-term experiments.

As a functional benchmark, we first used flow-cytometric DNA-content analysis to examine their recovery from exposure to hydroxyurea (HU), a compound that induces replication stress by depletion intracellular dNTP pools, thereby impairing DNA replication progression (Fig. 1A). At the time of HU removal, all three lines were enriched for cells with G1/early-S DNA content; by 6 h, their distributions had shifted towards higher DNA content, indicating cell cycle progression from G1/early-S to late-S and G2/M (Fig. 1B). At 8 h, the profiles diverged, with *HUS1*^+/−^ cultures showing a larger fraction of cells with G1 DNA content and a corresponding smaller G2/M fraction than *WT* cultures, a difference that persisted through 12 h (Fig. 1B, C and D). These distributions are consistent with earlier passage through G2/M and return to G1 in HUS1-deficient cells. Conversely, *RAD9*^+/−^ cultures contained a smaller G1 fraction than *WT* at 8 h. This difference persisted at 12 h and was accompanied by an increased G2/M fraction, indicating delayed late-cell-cycle progression (Fig. 1B, C and D). Thus, consistent with our previous observations (29), partial loss of HUS1 and RAD9 produced opposing cell-cycle responses during recovery from RS.

**Figure 1.**
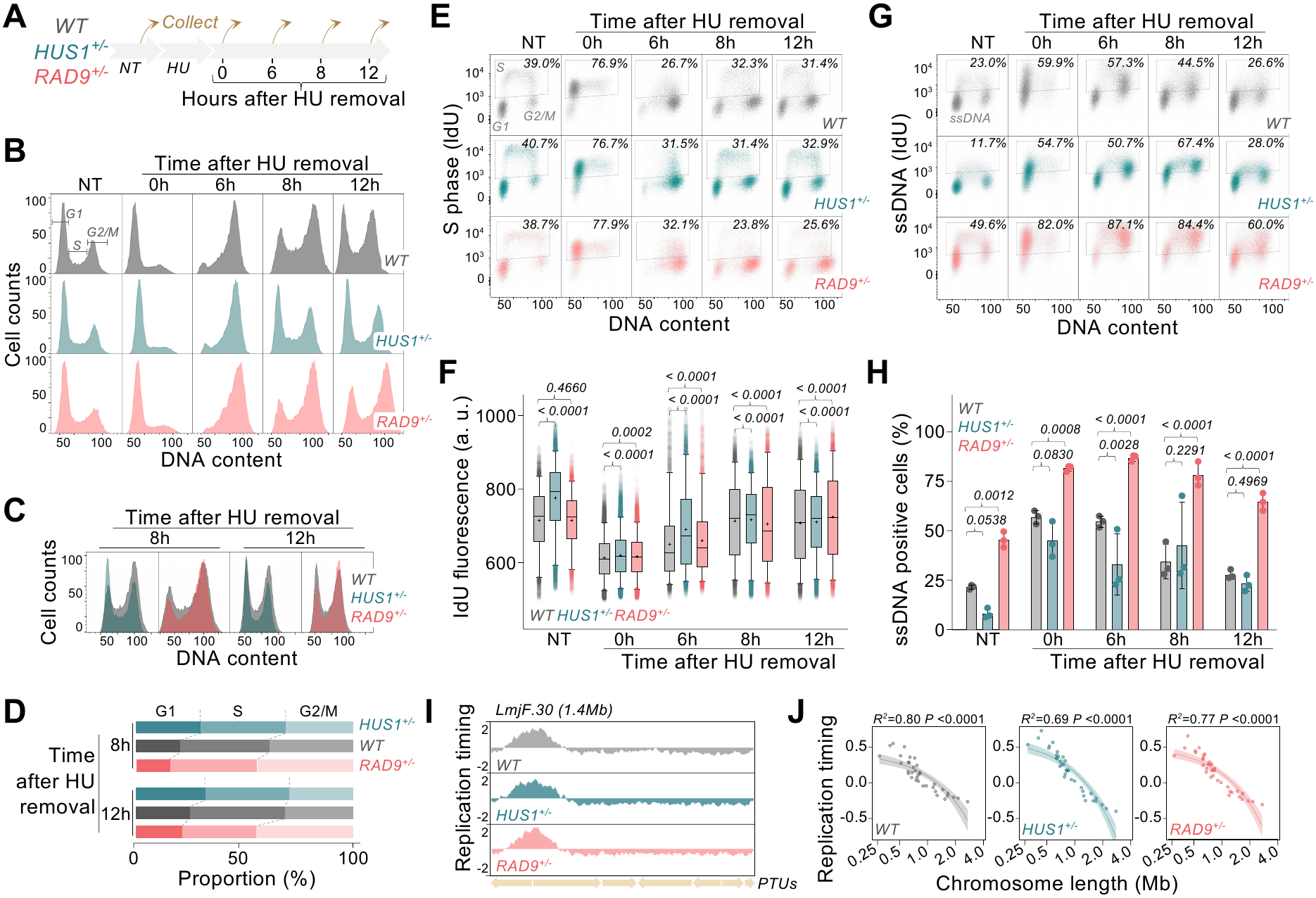
HUS1 and RAD9 deficiencies produce distinct DNA-replication and cell-cycle responses to replication stress without altering the basal replication-timing programme. **(A)** Experimental design. Wild-type (*WT*), *HUS1^+/−^* and *RAD9^+/−^ L. major* promastigotes were either left untreated (NT) or exposed to hydroxyurea (HU; 10 mM for 15 h). Following HU removal, cells were collected immediately (0 h) or after 6, 8 and 12 h. (B) Representative DNA-content histograms obtained by flow cytometry under untreated conditions and at the indicated times after HU removal. G1, S and G2/M gates, used for quantifications at C and D, are indicated in the untreated *WT* profile. (C) Overlay of the *WT*, *HUS1*^+/−^ and *RAD9*^+/−^ DNA-content profiles at 8 and 12 h after HU removal. (D) Relative proportions of cells assigned to the G1, S and G2/M gates at 8 and 12 h after HU removal. (E) Representative bivariate flow-cytometry plots of incorporated IdU fluorescence against DNA content. Dashed gates identify IdU-positive S-phase cells, and values indicate the percentage of cells within this gate. (F) Quantification of IdU fluorescence within the gated S-phase populations, providing a single-cell measure of DNA-synthesis activity. Boxes encompass the interquartile range, horizontal lines indicate medians, crosses indicate means and whiskers extend to 5 to 95 percentiles. *P* values were determined with Kruskal-Wallis test. a.u., arbitrary units. (G) Representative bivariate plots of IdU fluorescence against DNA content following IdU detection without DNA denaturation. Dashed gates define the ssDNA-positive population, with the corresponding percentage indicated in each plot. (H) Quantification of ssDNA-positive cells under untreated conditions and following HU removal. Bars represent means ± SEM, and points represent three independent biological replicates. *P* values were determined with ordinary one-way ANOVA test. (I) Representative marker frequency sequencing (MFA-seq) profiles across chromosome 30 (LmjF.30; 1.4 Mb). Replication-timing values are plotted across the chromosome, and arrows beneath the profiles indicate the positions and transcriptional orientations of polycistronic transcription units (PTUs). (J) Relationship between chromosome-average replication timing and chromosome length for all 36 chromosomes in each cell line. Each point represents one chromosome. Fitted regression curves and their confidence intervals are shown, together with the corresponding *R*² and *P* values.

Whether the contrasting cell-cycle phenotypes caused by HUS1 and RAD9 deficiency were accompanied by altered DNA synthesis remained unknown. We therefore performed bivariate flow-cytometric analysis of 5’-iodo-2’-deoxyuridine (IdU) incorporation and DNA content. Given IdU is a thymidine analogue that is incorporated into newly synthesised DNA, it allowed us to quantify both the fraction of cells actively synthesising DNA (IdU-positive cells, as detected under denaturing conditions; see methods; Fig. 1E) and IdU fluorescence within the gated population as a measure of DNA-synthesis activity per cell (Fig. 1F). The IdU-positive fraction was 39-41% in untreated cultures but increased to 77-78% immediately after HU treatment in all three lines, indicating that comparable proportions of *WT*, *HUS1^+/−^* and *RAD9^+/−^* cells were engaged in DNA synthesis at this time point. By 6 h, as the populations progressed towards G2/M (Fig. 1B), this fraction had declined in all three lines but remained modestly higher in both heterozygous mutants than in *WT* cells (Fig. 1E). At 8 and 12 h, the proportion of IdU-positive *HUS1*^+/−^ cells was similar to *WT,* although IdU fluorescence was significantly higher in *HUS1*^+/−^ cells under untreated conditions and at all time point after HU removal (Fig. 1F). A more dynamic phenotype was observed upon RAD9 deficiency. The IdU-positive fraction was comparable between *RAD9*^+/−^ and *WT* cells immediately after HU removal and was modestly higher in *RAD9*^+/−^ cells at 6 h but fell below that of *WT* cultures at both 8 and 12 h (Fig. 1E). IdU fluorescence was comparable between *RAD9^+/-^*and *WT* cells under untreated conditions but was significantly reduced at 8 h after HU removal (Fig. 1F). Together, these findings indicate that HUS1 and RAD9 deficiencies differentially affect DNA-synthesis dynamics during recovery from RS: HUS1 deficiency is associated with sustained higher IdU incorporation per DNA-synthesising cell, whereas RAD9 deficiency results in a transient reduction in IdU incorporation followed by a sustained decrease in the fraction of cells engaged in DNA synthesis.

To determine if and how HUS1 and RAD9 deficiency altered ssDNA accumulation, a hallmark of RS, we quantified IdU accessibility by flow cytometry (Fig 1G). Following IdU labelling, incorporated IdU was detected without prior DNA denaturation; under these conditions, antibody binding reports accessible IdU-containing ssDNA rather than total IdU incorporation (Fig. 1G and H). Compared with *WT* cells, *HUS1*^+/−^ cells showed a lower median ssDNA-positive fraction under untreated conditions, although this difference did not reach statistical significance. Following HU removal, a similar reduction was detected at 0 h, followed by a significant reduction at 6 h (Fig. 1H), when the population was progressing through late S phase towards G2/M (Fig. 1B). This transient reduction is consistent with decreased formation or persistence of detectable ssDNA intermediates in HUS1-deficient cells, although the assay cannot distinguish between reduced ssDNA exposure and more rapid processing. By contrast, *RAD9*^+/−^ cells displayed a significantly greater ssDNA-positive fraction than *WT* cells under untreated conditions and at every time point following HU removal (Fig. 1G, H). Notably, ssDNA remained elevated at 8 and 12 h despite the reduced proportion of *RAD9*^+/−^ cells actively synthesising DNA at these time points (Fig. 1E). These findings support a role for RAD9 in limiting ssDNA accumulation or promoting its resolution during both unperturbed growth and recovery from RS. Thus, HUS1 deficiency transiently reduced detectable ssDNA during late recovery, whereas RAD9 deficiency caused its sustained accumulation. These findings show that appropriate HUS1 and RAD9 dosage is required for coordinated recovery from RS in *L. major*, but that the two subunits make distinct contributions to this response. Some effects were already evident without HU treatment (namely, increased IdU incorporation in *HUS1*^+/−^ cells and elevated ssDNA in *RAD9*^+/−^ cells) indicating that both proteins also contribute to unperturbed DNA replication.

We next asked whether the RS phenotypes associated with HUS1 or RAD9 deficiency extended to the unperturbed genome-replication programme. Previous Marker Frequency Analysis (MFA-seq) revealed an unusual replication timing programme in *L. major* in which a single dominant early-replicating region is detected on each chromosome (32), generating an asymmetric size-dependent inter-chromosomal timing profiles, with larger chromosomes replicating, on average, later than smaller ones (Fig. 1I, J) (27, 30). We therefore generated MFA-seq profiles from *WT*, *HUS1*^+/−^ and *RAD9*^+/−^ cells grown in the absence of HU. The genome-wide profiles were highly similar among the three lines, with no detectable changes in the dominant early-replicating regions (Fig. 1I). Moreover, all three lines retained the strong chromosome-size-dependent replication timing pattern observed in *WT* cells (Fig. 1J). Thus, within the population-level resolution of MFA-seq, partial loss of HUS1 or RAD9 did not cause major reorganisation of the unperturbed replication-timing programme, despite profoundly affecting cell-cycle recovery, IdU incorporation and ssDNA accumulation following RS.

Collectively, these findings suggest that HUS1 and RAD9 appear to regulate the execution and processing of DNA synthesis in response to RS, rather than the global temporal organisation of chromosome duplication.

### MTX selection differentially alters HUS1 and RAD9 abundance without severely compromising parasite growth

Having established that partial loss of HUS1 or RAD9 produces distinct cell-cycle and DNA RS phenotypes, we next asked whether this functional divergence extends to adaptive genome plasticity. For this, *WT*, *HUS1*^+/−^ and *RAD9*^+/−^ populations were subjected to stepwise selection from 10 to 500 μM methotrexate (MTX), with samples collected after five passages at each concentration. Parallel selection series were performed with or without pre-exposure to HU to determine whether additional RS influenced the response to MTX (Fig. 2A).

**Figure 2.**
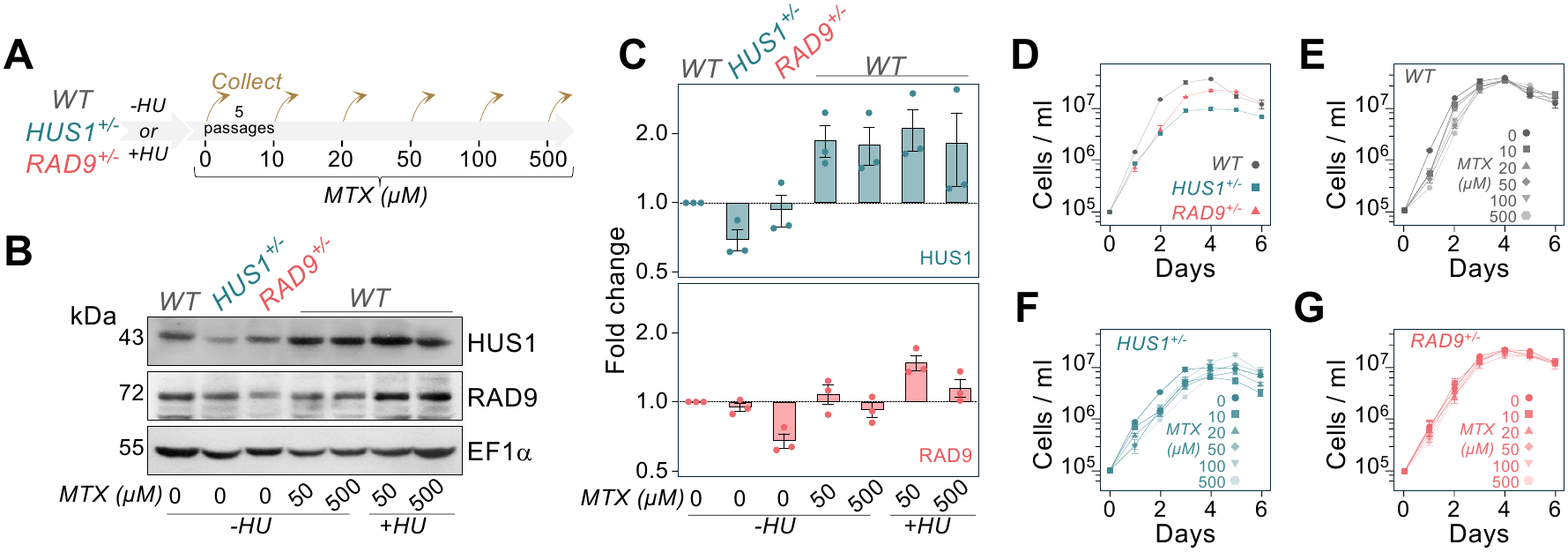
Stepwise MTX selection differentially modulates HUS1 and RAD9 abundance without causing major growth defects. **(A)** Experimental design for methotrexate (MTX) selection. *WT*, *HUS1*^+/−^ and *RAD9*^+/−^ *L. major* promastigotes were either left untreated (-HU) or pre-exposed to hydroxyurea (+HU; 10 mM for 15 h) before sequential selection with 10, 20, 50, 100 and 500 μM MTX. Cultures were maintained for five passages at each concentration before collection and progression to the next selection step. **(B)** Representative immunoblot analysis of HUS1 and RAD9 abundance. Unselected *WT*, *HUS1*^+/−^ and *RAD9*^+/−^ cells were compared with *WT* populations selected at 50 or 500 μM MTX with or without prior HU exposure. EF1α served as the loading control. **(C)** Densitometric quantification of HUS1 (upper panel) and RAD9 (lower panel) abundance. Signals were normalised to EF1α and expressed relative to unselected *WT* cells. Bars represent means ± SEM, and points represent independent biological replicates. **(D)** Growth profiles of unselected *WT*, *HUS1*^+/−^ and *RAD9*^+/−^ cells. **(E-G)** Growth profiles of *WT* (**E**), *HUS1*^+/−^ (**F**) and *RAD9*^+/−^ (**G**) populations selected and subsequently cultured at the indicated MTX concentrations. Cell densities were monitored for six days and are plotted on a logarithmic scale. Symbols and colour shades distinguish the MTX concentrations; values represent means ± SEM from three independent biological replicates.

Before examining the genomic outcomes of these selections, we assessed HUS1 and RAD9 abundance by immunoblotting. Analysis of unselected cells confirmed reduced abundance of the corresponding protein in *HUS1*^+/−^ and *RAD9*^+/−^ cells, whereas the reciprocal 9-1-1 subunit was comparatively little affected (Fig. 2B, C). MTX selection produced markedly different effects on the two proteins in *WT* cells. HUS1 abundance increased approximately 1.5-to 2-fold in populations selected at both 50 and 500 μM MTX, irrespective of HU pre-exposure. By contrast, RAD9 showed no comparable sustained changes. In HU-exposed populations only, RAD9 showed a more pronounced increase at 50 μM, but abundance approached basal levels by 500 μM MTX (Fig. 2B, C). Thus, HUS1 and RAD9 abundance does not change in concert during MTX selection, further supporting non-equivalent regulation of these 9-1-1 subunits.

We next determined whether the mutations or prolonged MTX selection imposed major proliferative costs that could confound subsequent analyses of genome plasticity. Among unselected populations, both *RAD9*^+/−^ and *HUS1^+/−^* cells reached consistently lower densities than *WT* cells, confirming their baseline growth defect (Fig. 2D) (29). Within each genotype, populations recovered at the successive MTX concentrations remained proliferative and displayed broadly similar growth trajectories to their respective unselected controls (Fig. 2E, F, G and Supplementary Fig. 1). Modest reductions were apparent at some of the higher MTX concentrations, particularly in *HUS1*^+/−^ cells subjected to MTX selection without (Fig. 2F) and with pre-exposure to HU (Supplementary Fig. 1), but no population underwent a major loss of proliferative capacity. These results establish that all selected populations broadly retained a similar proliferative capacity when compared to their unselected counterparts, and argue that subsequent differences in genome plasticity are unlikely to arise simply from severe growth impairment.

### RAD9 and HUS1 deficiencies differentially alter the extent and architecture of MTX-selected *DHFR-TS* amplification

MTX is known to select for amplification of loci containing genes associated with folate-metabolism in *Leishmania*, particularly those encoding dihydrofolate reductase-thymidylate synthase (*DHFR-TS*) and pteridine reductase 1 (*PTR1*) (33, 34). qPCR analysis detected no significant change in *PTR1* copy number following MTX selection in any of the three cell lines, irrespective of prior HU exposure (Supplementary Fig. 2), arguing against *PTR1* amplification as the principal resistance mechanism selected in these populations. By contrast, *DHFR-TS* copy number increased significantly across all three selected lines (Fig. 3A), identifying amplification of this locus as the predominant CNV associated with MTX adaptation.

**Figure 3.**
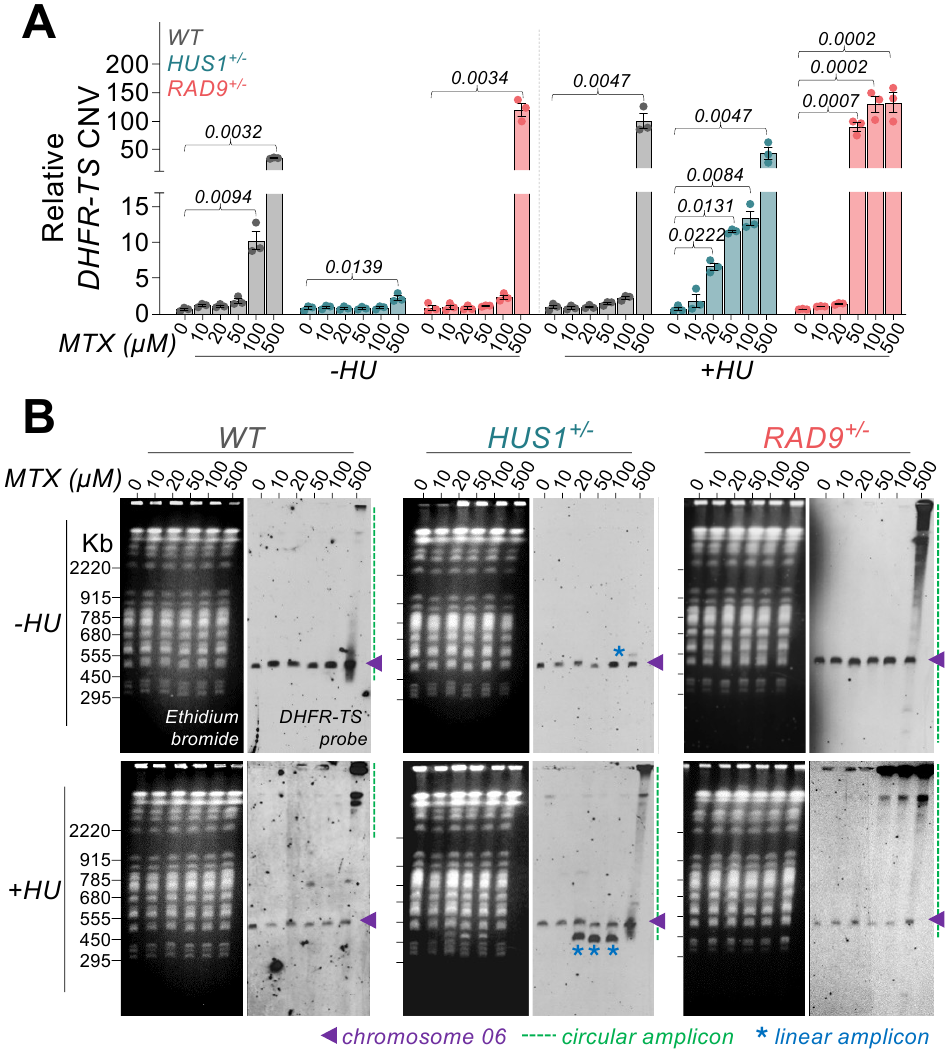
HUS1 and RAD9 deficiencies differentially alter the abundance and topology of MTX-selected *DHFR-TS* amplicons. *WT*, *HUS1*^+/−^ and *RAD9*^+/−^ *L. major* promastigotes were subjected to sequential selection with MTX, as outlined in Figure 2. **(A)** Quantitative PCR measurement of *DHFR-TS* copy number in MTX-selected populations with or without prior HU exposure. Ct values were normalised to G6PDH and expressed relative to Ct value from unselected *WT*. Bars represent means ± SEM from three independent biological replicates. *P* values were determined with Friedman multiple comparisons test. **(B)** Pulsed-field gel electrophoresis (PFGE) and Southern-blot analysis of the indicated MTX-selected populations. Intact chromosomes and extrachromosomal DNA molecules were visualised by ethidium-bromide staining prior to transferring. Membranes were hybridised with a *DHFR-TS* probe. Purple arrowheads, signal of intact chromosome 6. Blue asterisks, linear amplicons. Green dashed lines, circular amplicons.

We next assessed *DHFR-TS* amplification throughout the MTX selection series using two complementary approaches: qPCR and pulsed-field gel electrophoresis (PFGE) followed by Southern hybridisation (Fig. 3A, B). Whereas qPCR measures total locus dosage, PFGE-Southern analysis distinguishes the chromosome 6 *DHFR-TS* locus from extrachromosomal amplicons based on their topology-dependent migration. Linear amplicons resolve as discrete, size-defined bands, whereas circular molecules migrate anomalously, often producing diffuse signals or remaining trapped within or near the loading wells (35).

In populations selected without prior HU exposure, *WT* cells showed significant *DHFR-TS* amplification beginning at 100 µM MTX (Fig. 3A). However, a PFGE/Southern signal consistent with a circular extrachromosomal amplicon was detected only at 500 µM MTX (Fig. 3B), indicating that the increased *DHFR-TS* dosage at 100 µM preceded the accumulation of an extrachromosomal species detectable by this assay. By contrast, *HUS1*^+/−^ cells exhibited only a modest increase in *DHFR-TS* copy number at 500 µM MTX, accompanied by a single detectable linear amplicon. *RAD9*^+/−^ cells likewise showed significant amplification only at 500 µM MTX (Fig. 3A), but its magnitude was markedly greater and coincided with the accumulation of abundant circular amplicons (Fig. 3B). Thus, without prior HU exposure, the two deficiencies produced sharply contrasting outcomes at the highest MTX concentration: restricted amplification associated with a linear amplicon in *HUS1*^+/−^ cells, but extensive circular amplification in *RAD9*^+/−^ cells.

Prior HU exposure substantially altered both the dose dependence and topology of *DHFR-TS* amplification, particularly in the deficient lines. In WT cells, significant amplification and a PFGE/Southern signal consistent with a circular amplicon were detected only at 500 µM MTX (Fig. 3A, B). This signal appeared more abundant and heterogeneous than that observed in the corresponding population selected without prior HU exposure. In *HUS1*^+/−^ cells, however, *DHFR-TS* copy number increased significantly from 20 µM MTX onwards, with linear amplicons detected between 20 and 100 µM MTX (Fig. 3A, B). At 500 µM MTX, the linear species was no longer detectable. Instead, a signal consistent with a circular amplicon was detected at this MTX concentration. Reprobing the Southern blot with a telomeric probe showed that the amplicons detected between 20 and 100 µM MTX, but not at 500 µM, contained telomeric sequences, supporting a transition from linear to circular amplicon architecture during selection of *HUS1^+/-^*cells (Supplementary Fig. 3). In *RAD9*^+/−^ cells, the qPCR and PFGE/Southern results were closely concordant: pronounced *DHFR-TS* amplification and abundant circular amplicons were detected from 50 µM MTX onwards (Fig. 3A, B).

To determine whether the topology transition observed in *HUS1*^+/−^ cells depended on the nature of the preceding RS, we also examined *DHFR-TS* CNV following MTX selection after camptothecin (CPT) exposure. *WT* cells accumulated pronounced circular amplicons at 500 µM MTX, resembling the response following HU exposure, whereas *HUS1^+/−^* cells produced only linear amplicons from 50 µM MTX onwards (Supplementary Fig. 4). HU depletes dNTP pools and stalls replication forks, whereas CPT stabilises covalent topoisomerase I–DNA cleavage complexes that can be converted into double-strand breaks during replication. The contrasting amplicon profiles therefore suggest that the topology of *DHFR-TS* amplification in *HUS1^+/−^* cells is influenced by the nature of the preceding replication-associated lesion.

Together, these findings show that reduced HUS1 and RAD9 abundance differentially affects both the MTX concentration at which *DHFR-TS* amplification becomes detectable and the physical architecture of the resulting amplicons. HUS1 deficiency strongly limited amplification in the absence of prior RS but was associated predominantly with linear amplicons at low and intermediate MTX concentrations following HU exposure. Conversely, RAD9 deficiency was associated with extensive circular amplification, which emerged at lower MTX concentrations following HU exposure. These results therefore identify distinct contributions of HUS1 and RAD9 to the emergence or selection of locus-specific genome amplification under both normal and replication-stressed conditions.

### Genome-wide detection of circle-compatible rearrangements in *Leishmania*

We next asked whether HUS1 or RAD9 deficiency affects genome plasticity more broadly. Although circular and other amplicons have been characterised at individual *Leishmania* loci (36–39), dedicated circular or linear extra-chromosomal DNA enrichment workflows have not, to our knowledge, been benchmarked genome-wide in these parasites. We therefore adapted the Circle_finder junction-detection strategy (40) to paired-end bulk whole-genome sequencing. The algorithm identifies discordant read pairs containing split alignments that span non-reference end-to-start junctions compatible with circularisation. Because our libraries were not enriched for circular DNA, detection is biased towards abundant structures and likely underestimates rare or low-copy events. Nevertheless, the approach provides a genome-wide, locus-agnostic, readout of junction-bearing rearrangements. Representative calls on chromosomes 22 and 30 were supported by multiple high-quality split-read alignments converging on the same inferred boundaries (Fig. 4A).

**Figure 4.**
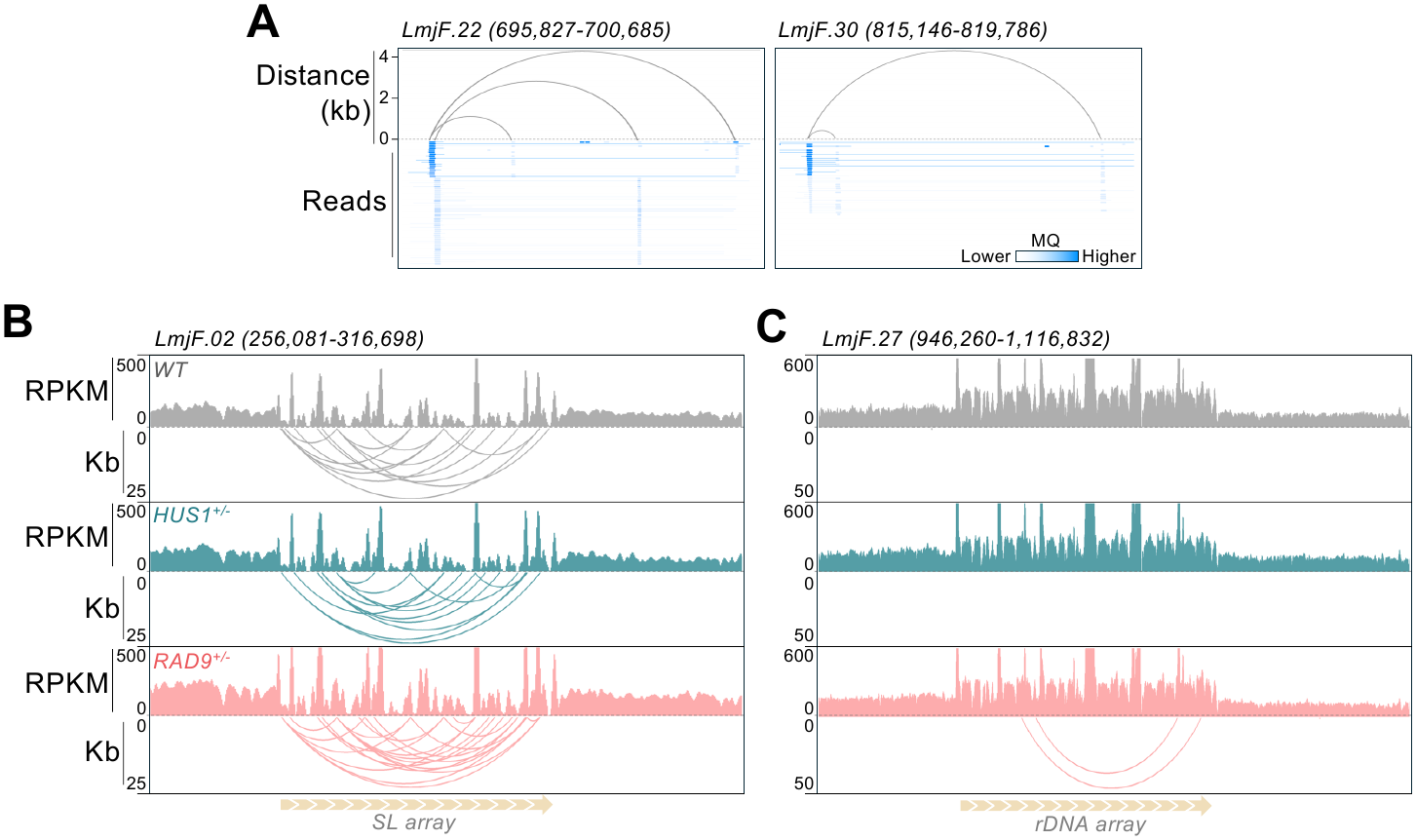
Breakpoint-supported detection of circle-compatible rearrangements from bulk *L. major* whole-genome sequencing data. **(A)** Representative *WT* calls at indicated loci. Arcs connect the inferred boundaries of each rearranged interval, with their height representing the genomic distance between junction endpoints. Read alignments supporting the corresponding non-reference junctions are shown beneath each candidate and coloured according to mapping quality (MQ). **(B)** and **(C)** Normalised sequencing coverage, shown as reads per kilobase per million mapped reads (RPKM), and junction evidence across the *SL* and rDNA arrays, respectively.

To assess the biological plausibility of these calls and whether repeat content alone drove their detection, we examined the spliced-leader (*SL*) RNA gene and ribosomal DNA (*rDNA*) arrays. Multiple rearrangement-compatible junctions were detected within the *SL* array in *WT*, *HUS1^+/−^*and *RAD9^+/−^* cells (Fig. 4B), consistent with its documented spontaneous expansion and recombination-associated remodelling (41, 42). By contrast, no junctions were detected within the similarly repetitive *rDNA* array in *WT* or *HUS1^+/−^* cells, whereas only two *rDNA*-associated junctions were identified in *RAD9^+/−^*cells (Fig. 4C). Although *rDNA* copy number and internal organisation vary among *Leishmania* isolates (43, 44), this sparse, line-restricted signal argues that repeat density alone did not generate widespread calls.

Because end-to-start junctions can also arise from tandem duplications and related rearrangements, they are not uniquely diagnostic of circular topology. We therefore refer to these calls as circle-compatible rearrangements. Within this scope, it provides a deliberately conservative but robust genome-wide readout of junction-bearing structural variation, enabling comparison of the effects of HUS1 and RAD9 deficiency beyond the *DHFR-TS* locus.

### HUS1 and RAD9 deficiencies differentially reshape the genome-wide rearrangement landscape

Next, we compared the genome-wide distribution of circle-compatible rearrangements among *WT*, *HUS1^+/−^*and *RAD9^+/−^* cells. Chromosome 36 illustrates the number and sizes of candidate events among the three lines (Fig. 5A). Genome-wide, events were detected on every chromosome but were unevenly distributed, with 818 calls in *WT*, 486 in *HUS1^+/−^*and 759 in *RAD9^+/−^* cells (Fig. 5B). Normalisation by chromosome length revealed distinct chromosome-specific profiles, indicating that these differences were not explained solely by chromosome size (Fig. 5C). Notably, *HUS1^+/−^*cells displayed lower event densities across most chromosomes than either *WT* or *RAD9^+/−^* cells. Thus, reduced HUS1 dosage is associated with a genome-wide decrease in detectable circle-compatible rearrangements, consistent with HUS1 contributing to their formation or retention within the population.

**Figure 5.**
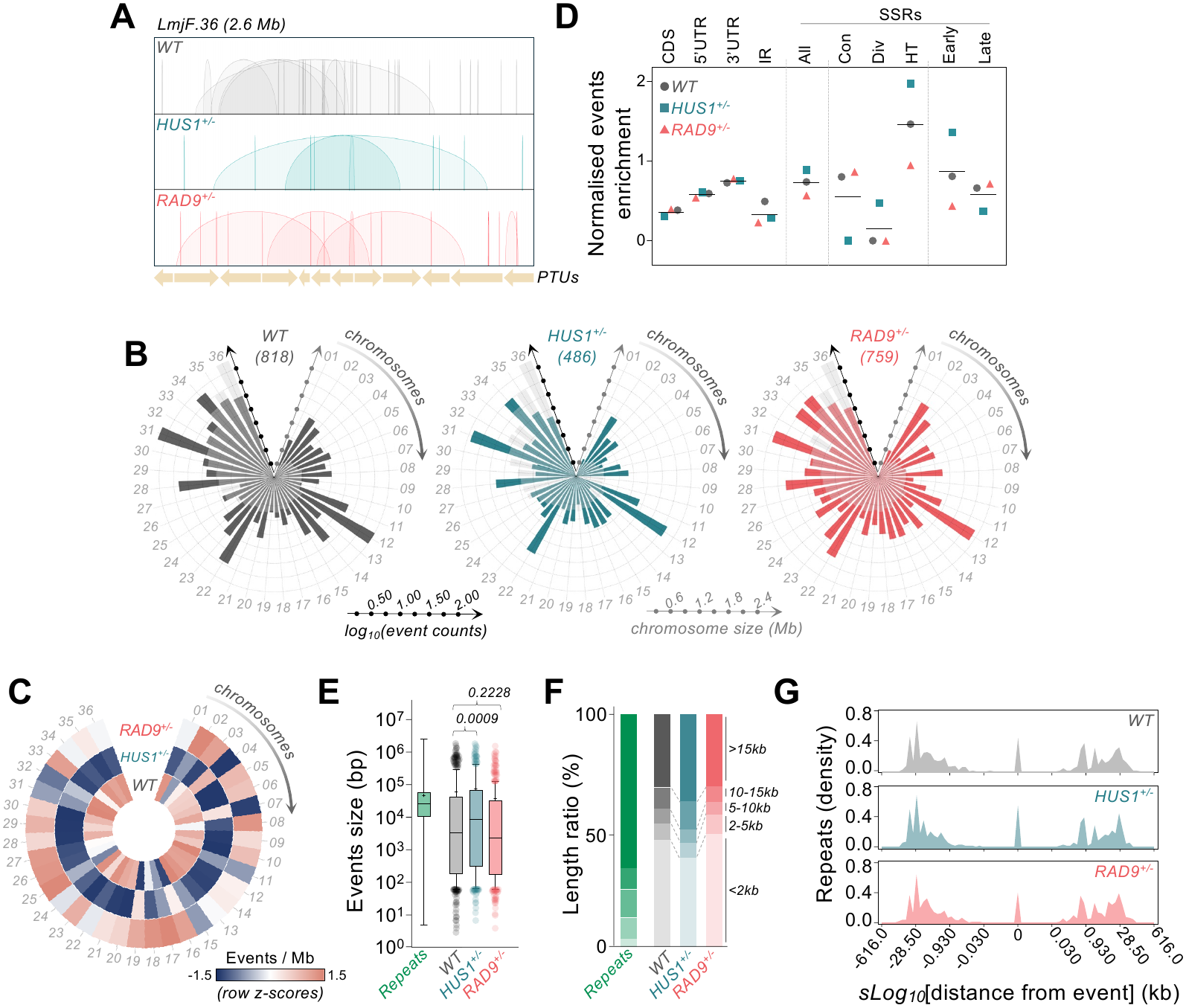
HUS1 and RAD9 deficiencies differentially reshape the genome-wide burden, distribution and size of circle-compatible rearrangements. **(A)** Distribution of circle-compatible rearrangements (arcs) across chromosome LmjF.36. Events too small to resolve as arcs at chromosome scale appear as near-vertical features. PTUs, polycistronic transcription units. **(B)** Radial pots showing the number of events in each chromosome. Coloured radial bars, log_10_-transformed number of events per chromosome. Light grey bars, chromosome length in megabases (Mb). Numbers in parentheses, genome-wide circle-compatible rearrangements counts. **(C)** Density of circle-compatible rearrangements per chromosome. Values are displayed as row-wise z-scores calculated independently for each genotype. **(D)** Normalised circle-compatible rearrangements enrichment calculated by dividing the percentage of all calls located within that feature by the percentage of the genome covered by the feature. CDS, coding sequences. UTR, untranslated regions. IR, intergenic regions. SSRs, strand-switch regions, were grouped into six categories: Con, convergent; Div, divergent; HT, head-to-tail; Early, early-replicating and Late, late-replicating. All, every SSR type combined. **(E)** Size distributions of repeat-predicted amplicons and circle-compatible rearrangements in the three cell lines. Repeat-predicted sizes were derived from the genomic distances between annotated repeat pairs. Boxes encompass the interquartile range, central lines indicate medians, whiskers extend from 5 to 95 percentiles. *P* values were determined with Kruskal-Wallis test. **(F)** Proportional distribution of repeat-predicted intervals and detected events among the indicated size classes. **(G)** Normalised density of annotated repeats surrounding the start coordinates of candidate events. For better resolution of density peaks, distances are displayed on a signed logarithmic scale (*sLog_10_*), with negative and positive values indicating repeats upstream and downstream of the event boundary, respectively, and zero denoting the event start position.

We next examined the genomic features associated with these events. After correcting for the proportion of the genome occupied by each feature, event representation within coding sequences (CDSs), 5′ and 3′ untranslated regions (UTRs) and intergenic regions (IRs) was broadly comparable among the three lines. At the boundaries between PTUs, however, event distribution varied according to boundary class, replication timing and genotype. Head-to-tail (HT) SSRs mark the termination of one PTU and initiation of the next, with both transcribed in the same direction; convergent SSRs separate PTUs transcribed towards one another and are associated with transcription termination; and divergent SSRs separate PTUs transcribed away from one another and are associated with transcription initiation. SSRs were also classified as early- or late-replicating according to when they are duplicated during S phase. HT and early-replicating SSRs showed the strongest event enrichment in all three lines, with the highest values in *HUS1*^+/−^ cells and the lowest in *RAD9*^+/−^ cells relative to *WT* (Fig. 5D). Conversely, *HUS1*^+/−^ cells showed particularly low enrichment at convergent and late-replicating SSRs, but greater enrichment at divergent SSRs than either *WT* or *RAD9*^+/−^ cells (Fig. 5D). These non-uniform distributions indicate that detectable rearrangements preferentially associate with specific genomic environments and that HUS1 and RAD9 deficiencies reshape these associations differently.

Candidate-event sizes spanned several orders of magnitude and broadly overlapped the range of amplicon sizes predicted from genomic repeat pairs (Fig. 5E) (18). Events detected in *HUS1^+/−^* cells were significantly larger than those in *WT*, whereas the size distribution in *RAD9^+/−^*did not differ significantly from *WT*. Comparison of observed and repeat-predicted size-class distributions showed that most potential repeat-defined amplicons were larger than 15 kb, whereas most detected events in all three lines were smaller than 15 kb (Fig. 5F). This discrepancy could reflect preferential detection of shorter structures by our approach or indicate that only a subset of repeat-defined rearrangements arises or persists at detectable abundance in vivo. The size-class distributions were broadly comparable between *WT* and *RAD9^+/−^*, whereas *HUS1^+/−^* cells displayed a smaller proportion of events below 2 kb and a corresponding increase in the classes above 10 kb (Fig. 5F). Together with the lower total number of calls, this shift suggests that HUS1 deficiency preferentially reduces the generation or population-level retention of smaller rearrangements, an effect not observed following RAD9 deficiency.

Repeat-density analysis revealed a prominent peak at the inferred start position of candidate rearrangements, accompanied by secondary peaks approximately 0.9 and 30 kb at both upstream and downstream (Fig. 5G). This distribution is compatible with repeat-associated rearrangement boundaries across the size range represented by both the predicted and detected events (Fig. 5E, F). These repeat-density profiles, as well as sequencing depth across repeats (Supplementary Fig. 5), were nevertheless broadly similar among *WT*, *HUS1^+/−^*and *RAD9^+/−^* cells, indicating that their differences in event burden and size distribution were unlikely to be explained by a major shift in repeat proximity or repeat copy number.

Together, these analyses reveal distinct consequences of the two deficiencies: HUS1 deficiency was associated with fewer detectable events but a shift towards larger rearrangements and specific genomic contexts, whereas RAD9 deficiency retained a largely *WT*-like overall event number and size distribution. Thus, RAD9 and HUS1 make non-equivalent contributions to the genome-wide control of structural plasticity in *L. major*.

### HUS1 and RAD9 differ in repeat association and locus-specific CNV phenotypes

Because repeated sequences are major substrates for *Leishmania* genome rearrangements (18), we next asked whether the distinct CNV phenotypes associated with HUS1 and RAD9 deficiency might reflect different interactions with repetitive DNA. We used our previously reported ChIP-seq analysis (45) to compare HUS1 and RAD9 deposition profiles around annotated repeats under untreated and HU-exposed conditions. RPA1, an ssDNA-binding component of the replication-stress response (29, 46), and γH2A, a marker of DNA damage and RS in *Leishmania* (29, 31, 45, 47), were included for comparison (Fig. 6A). All four ChIP-seq signals displayed structured profiles around the repeats that were absent from genome-shuffled controls, indicating non-random associations. However, their spatial distributions differed markedly: HUS1 formed two peaks flanking the repeats, whereas RAD9 and RPA1 showed broader repeat-centred accumulation and γH2A displayed a narrower central peak. HU exposure caused only modest changes in signal intensity and largely preserved these distinct profiles. These profiles suggest that repetitive sequences constitute endogenous sites of replication-associated genome surveillance in *L. major*, with HUS1, RAD9 and RPA1 displaying distinct patterns of association at these regions rather than being recruited solely in response to acute RS. Thus, although RAD9 and HUS1 are canonical subunits of the same 9-1-1 checkpoint clamp, their distinct repeat-association profiles may contribute to their non-equivalent effects on *DHFR-TS* amplification and genome-wide rearrangement formation.

**Figure 6.**
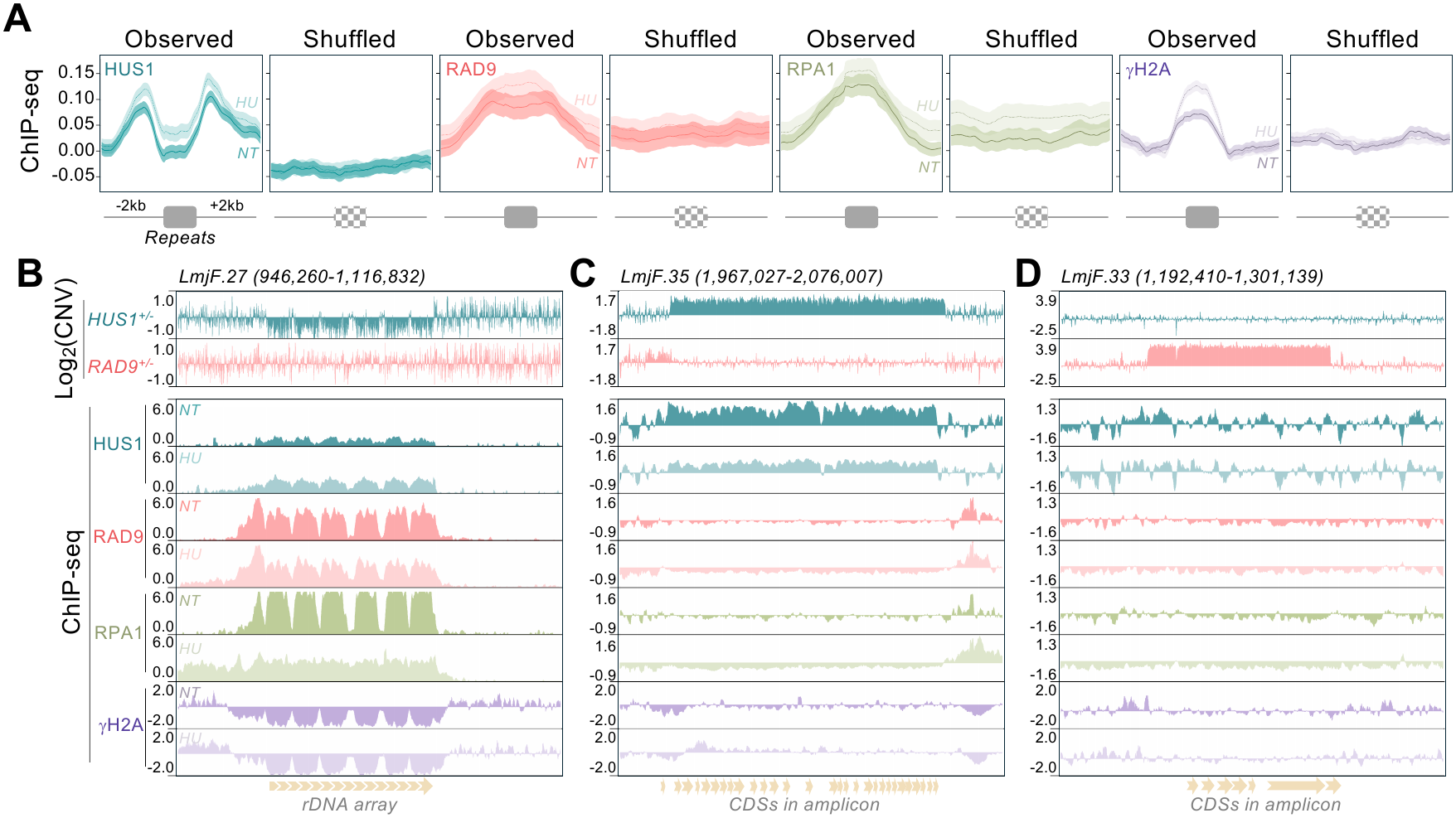
HUS1 and RAD9 display distinct repeat-association profiles and non-equivalent locus-specific CNV phenotypes. **(A)** Metaplots of normalised ChIP-seq signal for HUS1, RAD9 and RPA1, together with the DNA-damage-associated histone mark γH2A. Profiles are shown across ± 2 kb windows centred on annotated repeats (observed) as compared to genome-shuffled repeat coordinates (shuffled). Darker- and lighter-coloured profiles represent ChIP-seq signal from untreated (NT) and HU-treated (5 mM, 8 h) cells, respectively; shaded areas around lines indicate SEM. **(B - D)** Genome-browser views comparing locus-specific copy-number variation with ChIP-seq profiles. Upper tracks show the log₂-transformed copy-number ratio in *HUS1*+/− and *RAD9*+/− cells relative to *WT*. Positive and negative values indicate increased and decreased sequencing reads abundance, respectively. Lower tracks show normalised ChIP-seq signal under NT and HU-exposed conditions.

We then integrated the ChIP-seq profiles with genotype-specific CNVs at three representative loci (Fig. 6B–D). At the chromosome 27 *rDNA* array, *HUS1^+/−^* cells displayed a broad reduction in sequence coverage, consistent with reduced *rDNA* copy number, whereas no comparable change occurred in *RAD9^+/−^* cells (Fig. 6B). The occupancy profiles at this locus were also distinct: under untreated conditions, RAD9 and RPA1 showed pronounced accumulation across the array, whereas HUS1 displayed a more modest signal. Following HU exposure, HUS1 accumulation increased slightly, while the RAD9 and RPA1 profiles decreased or became more evenly distributed. By contrast, γH2A was relatively depleted across the array under both conditions. Thus, *rDNA* copy-number maintenance was selectively sensitive to HUS1 deficiency despite the association of several replication-stress-response factors with this locus.

A different relationship was observed at a chromosome 35 locus that underwent broad copy-number gain specifically in *HUS1^+/−^*cells (Fig. 6C). HUS1 accumulated throughout this region under untreated conditions, and this association was attenuated following HU exposure. Neither RAD9, RPA1 nor γH2A showed comparable interval-wide accumulation, although focal RAD9 and RPA1 signals were apparent near one boundary. The spatial correspondence between HUS1 occupancy and the HUS1-dependent CNV raises the possibility that HUS1 contributes directly to copy-number control at this locus.

Conversely, a locus at chromosome 33 was strongly amplified in *RAD9^+/−^*cells but remained unchanged in *HUS1^+/−^* cells (Fig. 6D). None of the four factors showed substantial accumulation across this region under either untreated or HU-exposed conditions. Thus, this locus shows that the CNV increase associated with RAD9 deficiency need not coincide with detectable local occupancy by RAD9, HUS1 or RPA1, or with γH2A enrichment.

Together, these findings reveal distinct modes of HUS1 and RAD9 association with repetitive DNA and a locus-dependent relationship between replication-stress-factor occupancy and CNV. Some genotype-specific alterations coincide with local factor accumulation, whereas others arise without detectable enrichment, indicating that occupancy alone does not predict susceptibility to copy-number change. Instead, HUS1 and RAD9 appear to influence genome plasticity through functionally compartmentalised mechanisms shaped by local genomic context.

### RAD9 deficiency is associated with broader chromosome-somy changes than HUS1 deficiency

Because *Leishmania* genome plasticity encompasses both local rearrangements and chromosome-copy-number variation, we next examined whether HUS1 and RAD9 deficiencies also altered chromosome somy. We inferred population-average somy for all 36 chromosomes from normalised whole-genome sequencing read depth. Although the overall somy profiles were broadly conserved, discrete genotype-dependent differences were evident (Fig. 7A). Plotting somy fold changes relative to *WT* against chromosome length revealed no consistent size dependence: *HUS1^+/−^* cells displayed comparatively restricted changes, whereas *RAD9^+/−^* cells exhibited broader changes (Fig. 7B).

**Figure 7.**
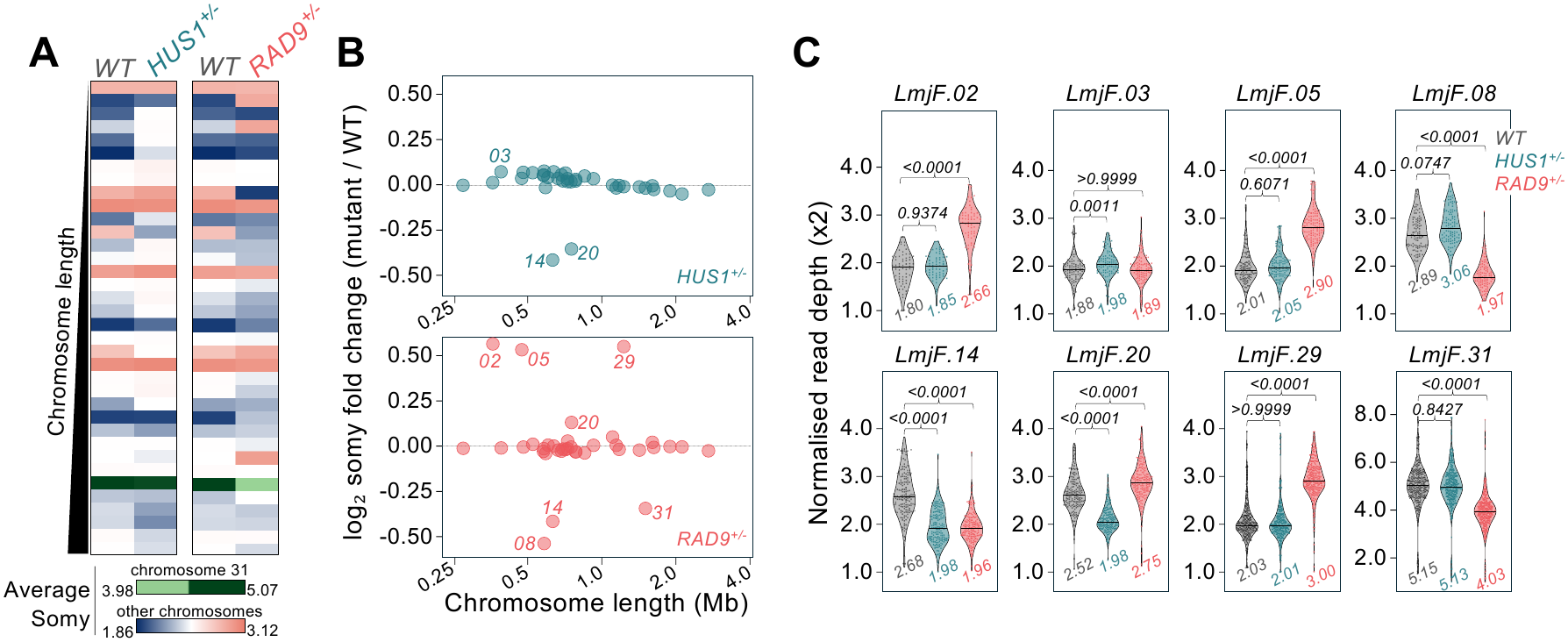
Population-average chromosome-somy profiles in *WT*, *HUS1*+/− and *RAD9*+/− *L. major*. **(A)** Colourmaps of chromosome somy inferred from normalised whole-genome sequencing read depth. Each row represents one of the 36 chromosomes, with relative chromosome length indicated on the left. For better comparison, *WT* is shown alongside *HUS1*+/− or *RAD9*+/−. Chromosome 31 is displayed using a separate green scale because of its higher somy. **(B)** Log₂-transformed somy ratios for *HUS1*+/− (upper panel) and *RAD9*+/− (lower panel) relative to *WT*, plotted against chromosome length. Chromosomes with copy number changes are labelled. **(C)** Distributions of normalised read depth across the indicated chromosomes. Values beneath each violin indicate the estimated population-average somy. *P* values were determined with Kruskal-Wallis test.

Because chromosome mosaic aneuploidy is pervasive in *Leishmania*, non-integer chromosome-wide somy estimates provide population-level evidence for the coexistence of cells carrying different chromosome-copy-number states. Although bulk sequencing cannot resolve their precise proportions or single-cell karyotypes, changes in average somy reveal redistribution of these states within the population. In *WT* cells, chromosomes 14 and 20 had average somies of 2.68 and 2.52, respectively, consistent with mosaic disomy and trisomy. Both decreased to 1.98 in *HUS1^+/−^* cells, indicating a marked shift towards populations dominated by disomic cells (Fig. 7C). Chromosome 3 also changed from 1.88 to 1.98; although statistically significant, this difference was modest, with both populations remaining predominantly disomic.

By contrast, *RAD9^+/−^* cells displayed extensive remodelling of chromosome somy. Chromosome 2 increased from 1.80 in *WT* to 2.66 in *RAD9^+/−^* cells, indicating a substantial redistribution towards higher-somy, most likely trisomic, cells. Chromosomes 5 and 29 similarly increased from near-disomic averages of 2.01 and 2.03 to near-trisomic averages of 2.90 and 3.00, respectively. Chromosome 20 increased more modestly from 2.52 to 2.75, consistent with greater representation of trisomic cells. Conversely, chromosomes 8 and 14 decreased from 2.89 and 2.68 to 1.97 and 1.96, respectively, indicating shifts towards populations dominated by disomic cells. The high-somy chromosome 31 also decreased from an average of 5.15 copies in *WT* to 4.03 in *RAD9^+/−^* cells, corresponding to a population-level shift from near-pentasomy to near-tetrasomy (Fig. 7C).

Altogether, HUS1 deficiency was associated with comparatively restricted shifts towards predominant disomy, whereas RAD9 deficiency produced broader, bidirectional redistribution of chromosome-somy states. Together with their distinct effects on local CNV and genome-wide rearrangements, these findings reveal non-equivalent roles for HUS1 and RAD9 in shaping both local and chromosome-scale genome plasticity in *L. major*.

## Discussion

Here, we combined genetic perturbation with targeted and genome-wide analyses to determine how reduced HUS1 or RAD9 abundance affects genome plasticity in *L. major*. Although both proteins are subunits of the 9-1-1 clamp, their deficiencies produced non-equivalent effects on DNA synthesis, ssDNA accumulation, extrachromosomal amplification, junction-bearing rearrangements and chromosome somy. These findings extend their established roles in DNA-damage responses, telomere homeostasis and genome maintenance (29, 31, 46). More broadly, they place RS surveillance at the interface between genome maintenance and potentially adaptive variation. Rather than acting simply as a general suppressor of genome variation, HUS1 and RAD9 appear to influence the types and genomic contexts of variants that arise or persist.

The divergent responses to HU further establish that HUS1 and RAD9 make distinct contributions to recovery from RS (29). *HUS1*^+/−^ cells showed earlier return to a G1-like DNA-content state, higher IdU incorporation per S-phase cell and transiently lower detectable ssDNA, whereas *RAD9*^+/−^ cells progressed more slowly and accumulated ssDNA persistently, including at later time points when fewer cells were synthesising DNA. Although IdU intensity is not a direct measure of fork speed, the combined measurements indicate that HUS1 and RAD9 deficiencies differentially affect the coordination of DNA synthesis and processing of RS intermediates. Their distinct γH2A responses after HU, as previously reported (29), are consistent with this interpretation and further indicate that the two deficiencies elicit distinct damage-associated chromatin responses during RS recovery.

An underlying basis for this divergence may lie in the unusual organisation of the 9-1-1 subunits in *Leishmania*. Size-exclusion chromatography recovered RAD1 predominantly in fractions compatible with the canonical clamp, whereas HUS1 and RAD9 were also detected in distinct molecular-mass fractions (29). Although these observations do not establish independent functions for alternative complexes, they provide a plausible molecular basis for the functional divergence observed here. Reduced abundance of either protein could therefore perturb canonical 9-1-1 activity while also affecting subunit-specific assemblies or interactions. This model accommodates their opposing phenotypes without requiring HUS1 and RAD9 to act antagonistically.

Despite the pronounced RS phenotypes, neither deficiency detectably reorganised the basal replication-timing programme in population-averaged MFA-seq. The dominant early-replicating locus on each chromosome and the chromosome-size-dependent timing relationship were preserved, indicating that changes in genome plasticity observed in HUS1- and RAD9-deficient cells are not explained by a global disruption of replication timing. Importantly, cell-cycle-resolved MFA-seq showed that HUS1 and RAD9 deficiencies affect replication timing at subtelomeres, whose replication extends beyond the conventional S phase window (30). This raises the possibility that some phenotypes observed here, including altered genome plasticity, may arise partly from changes in late subtelomeric replication. This is consistent with accumulating evidence linking the genome replication timing to genome variation in *Leishmania*. RAD51, which promotes repeat-mediated circular amplification (18), also helps maintain the normal temporal separation between the major early-replicating locus of each chromosome and its later-replicating subtelomeres (48). Loss of RNase H1 abolishes chromosome-size-dependent replication timing and alters aneuploidy and chromosome stability (27), whereas deletion of the ATR C-terminal region disrupts this timing relationship and changes the accumulation of indels and chromosome-specific CNVs (28). Together, these findings indicate that genome plasticity can be modulated through several features of the replication programme, not all of which are resolved by bulk timing profiles.

The contrasting *DHFR-TS* CNV phenotypes indicate that reduced HUS1 or RAD9 abundance and prior RS influence both the extent of amplification and the topology of the resulting extrachromosomal amplicons. Without prior HU exposure, HUS1 deficiency restricted circular amplification, whereas RAD9 deficiency ultimately yielded abundant circles. HU pre-exposure enabled *DHFR-TS* CNV to be detected at lower MTX concentrations in both mutants and was associated with genotype-specific amplification patterns. These phenotypes can be compared with established pathways of gene-amplification in *Leishmania*. RAD51 and RAD51-4 promote circular amplification (18, 49), and the restricted accumulation of *DHFR-TS* circles in *HUS1*^+/−^ cells phenotypically resembles the effect of RAD51 loss. By contrast, MRE11 nuclease activity supports efficient linear-amplicon formation (50), whereas the increased accumulation of circular amplicons in *RAD9*^+/−^ cells parallels the effect of MRE11 deletion. Although these phenotypic similarities do not establish that HUS1 or RAD9 participates directly in homologous recombination in *Leishmania*, the 9-1-1 clamp promotes

DNA-end resection and resection-dependent homologous recombination in other eukaryotes (51, 52). HUS1 and RAD9 may therefore differentially modulate the formation or persistence of ssDNA intermediates during repeat-mediated recombination, thereby influencing the production of circular and linear amplicons. This model does not require HUS1 and RAD9 to act antagonistically or either protein to oppose homologous recombination. Instead, they may cooperate with recombination factors to coordinate the timely processing or resolution of these intermediates, consistent with the opposing ssDNA-accumulation phenotypes caused by their respective deficiencies. However, because our assays measure accumulated amplicons, effects on their replication, inheritance or selective enrichment after formation remain plausible alternatives.

ChIP-seq analysis placed these effects within their chromosomal context. HUS1, RAD9, RPA1 and γH2A displayed distinct, non-random profiles around repeats in untreated cells that changed only modestly following HU exposure, suggesting that repetitive sequences are constitutively associated with replication-stress surveillance. Nevertheless, HUS1 and RAD9 exhibited markedly different spatial distributions: HUS1 concentrated at repeat flanks, whereas RAD9 showed broader enrichment extending across the repeats. These patterns may reflect engagement with distinct DNA intermediates, successive processing steps or alternative protein assemblies, although ChIP-seq cannot resolve the underlying DNA structures or establish direct activity at individual rearrangement boundaries.

Genome-wide, *HUS1*^+/−^ cells carried fewer but larger detectable circle-compatible junctions. By contrast, *RAD9*^+/−^ cells retained an event burden and size distribution broadly similar to WT, arguing against a general increase in rearrangement abundance despite their pronounced *DHFR-TS* phenotype. Notably, early-replicating SSRs, which overlap the major early-replicating locus on each chromosome, and HT SSRs, which mark transitions between transcription termination and initiation, showed the greatest event enrichment across all three lines. This distribution links rearrangement susceptibility to genomic regions with distinctive replication and transcriptional features. This is consistent with recent genome-wide mapping showing pronounced accumulation of HUS1, RAD9 and RPA1 at these regions (45). Together, these findings place rearrangement susceptibility at the intersection of replication and transcriptional organisation, while the opposing enrichment profiles in *HUS1*^+/−^ and *RAD9*^+/−^ cells support non-equivalent and context-dependent contributions of the two factors to rearrangement formation.

Comparing RAD9 and HUS1 occupancy with locus-specific CNV in their respective deficient lines further illustrates this context dependence. Reduced sequencing coverage across the *rDNA* array in *HUS1*^+/−^ cells suggests that HUS1 contributes to its maintenance. In yeast, the high transcriptional activity, repetitive organisation and specialised replication of *rDNA* arrays make them particularly dependent on recombination and RS control for copy-number homeostasis (53). Although this provides a useful mechanistic analogy, the pathways controlling *rDNA* dosage in *Leishmania* remain unresolved. Moreover, RAD9 occupancy across the same array exceeded that of HUS1, showing that local factor accumulation alone does not predict how its deficiency will affect copy number. Similarly, the amplified chromosome 33 locus in *RAD9^+/−^* cells showed no pronounced accumulation of RAD9, HUS1, RPA1 or γH2A. The clearest correspondence occurred at the chromosome 35 locus amplified in *HUS1*^+/−^ cells, where HUS1 accumulated strongly in *WT* cells. Local factor occupancy may therefore identify some vulnerable genomic contexts but is neither necessary nor sufficient to predict the direction or genotype specificity of CNV.

Somy analysis extends the consequences of 9-1-1 deficiency from local rearrangements to whole-chromosome copy-number variation. Whereas *HUS1*^+/−^ cells displayed changes in relatively few chromosomes, including shifts in population-average somy towards two copies, *RAD9*^+/−^ cells showed broader gains and losses. These population-level measurements must be interpreted in the context of *Leishmania* mosaic aneuploidy, in which the relative abundance of distinct somy states can change during passage and environmental transitions (21, 54, 55). Bulk sequencing therefore cannot distinguish the generation of new karyotypes from selection among pre-existing cells. Persistent ssDNA and delayed cell-cycle recovery in *RAD9*^+/−^ cells provide a plausible route to increased karyotypic variation, as under-replicated DNA entering division can promote chromosome breakage and segregation errors in other eukaryotes (56, 57). Whether related mechanisms connect RAD9 or HUS1 deficiency to somy changes in *Leishmania* remains unresolved. Notably, the lack of a consistent relationship between somy changes and chromosome length argues against a simple model in which the largest chromosomes are preferentially affected because they fail to complete replication.

In summary, our findings establish the 9-1-1 clamp, a replication stress response factor, as a determinant of genome plasticity in *L. major*. Reduced HUS1 and RAD9 abundance produces distinct patterns of genome variation, spanning locus-specific amplification, genome-wide junction landscapes and mosaic aneuploidy. Because viable heterozygotes were necessary for the long-term assays, longitudinal single-cell sequencing, together with topology-resolving enrichment of extra-chromosomal molecules, will be required to distinguish variant formation from subsequent selection and to determine the structures of the circle-compatible rearrangements. Despite these limitations, our data argue that the 9-1-1 response is not simply an impediment to genome variation; instead, our findings place the replication stress response pathway as a novel modulator of *Leishmania*’s genome plasticity.

## Material and Methods

### Parasite culture and cell lines generation

all cell lines were cultured as promastigotes at 26 °C in M199 medium supplemented with 40 mM HEPES, 0.1 mM adenine, 50 μg mL^-1^ biotin, 250 μg mL^-1^ haemin and 10% foetal bovine serum. *RAD9*^+/-^ and *HUS1*^+/-^ cell lines were generated from wild-type *Leishmania major* LT252 (MHOM/IR/84/LT252) as previously described (29, 46) and were selected with 40 μg mL^-1^ nourseothricin and 16 μg mL^-1^ hygromycin, respectively. MTX-resistant lines derived from wild-type, *RAD9*^+/-^ and *HUS1*^+/-^ cell lines mutants were maintained under the corresponding antibiotic selection described above, with MTX added at the concentrations used for their selection.

### Generation of MTX-resistant lines

MTX-resistant lines were generated by stepwise selection with increasing MTX concentrations (58). Cells were seeded at 1 x 10⁵ cells mL⁻¹ in MTX-containing medium and allowed to proliferate to approximately 1 x 10⁷ cells mL⁻¹ before being reseeded at 1 x 10⁵ cells mL⁻¹ in fresh medium containing the same MTX concentration. After five passages at each concentration, cells were transferred to the next higher MTX concentration.

### Preparation of whole-cell protein extracts and western blotting

Cells were harvested by centrifugation at 1,400 x *g* for 5 min, washed with 1 x PBS and resuspended in Laemmli sample buffer [50 mM Tris pH 6.5, 10 mM DTT, 0.1% (w/v) bromophenol blue and 10% (w/v) glycerol] at a density of 1 x 10⁵ cells μL⁻¹. Extracts were heated at 90 °C for 10 min, and proteins were resolved by SDS–PAGE on 10% polyacrylamide gels before transfer onto PVDF membranes with a pore size of 0.45 μm. Membranes were blocked for 1 h at room temperature in PBS-T [1× PBS containing 0.1% (v/v) Tween-20] supplemented with 10% (w/v) skimmed milk powder, then incubated for 2 h with primary antibodies diluted in PBS-T containing 3% (w/v) skimmed milk powder. Following four washes in PBS-T for 5 min each, membranes were incubated for 2 h with horseradish peroxidase-conjugated secondary antibodies diluted in PBS-T containing 3% (w/v) skimmed milk powder. Membranes were washed a further four times in PBS-T for 5 min each and incubated with ECL Prime Western Blotting Detection Reagent (Amersham). Signals were detected by exposure to Hyperfilm ECL (Amersham).

### Antibodies

Purified rabbit anti-HUS1 and anti-RAD9 antibodies were used at dilutions of 1:1,000 and 1:2,000, respectively (29, 46). Mouse anti-EF1α antibody (Roche) was used at 1:30,000. Horseradish peroxidase-conjugated secondary anti-rabbit secondary antibody (KPL, Gaithersburg, MD, USA) and anti-mouse secondary antibody (GE Healthcare, UK) were used at dilutions of 1:30,000.

### Pulsed-field gel electrophoresis (PFGE) and Southern blotting

Cells were harvested by centrifugation at 1,400 x *g* for 5 min and resuspended at 4 x 10⁸ cells mL⁻¹ in 2 x M199 medium without foetal bovine serum. The suspension was mixed with an equal volume of chromosome preparation buffer [130 mM NaCl, 142 mM Na₂HPO₄, 8 mM NaH₂PO₄ and 2% (w/v) glucose] containing 2% (w/v) low-melting-point agarose. The mixture was dispensed into acrylic moulds and incubated on ice for 20 min to allow solidification. Agarose plugs were incubated in lysis buffer [0.5 M EDTA, pH 9.0, 1% (w/v) SDS and 0.5 mg mL⁻¹ proteinase K] at 50 °C for 48 h. Plugs were subsequently washed in TE buffer (200 mM Tris-HCl and 100 mM EDTA) with constant agitation, replacing the buffer every 12 h.

Chromosomes were resolved by PFGE in 1% (w/v) agarose gels in 0.5 x TBE buffer (45 mM Tris, 45 mM boric acid and 1 mM EDTA) using a CHEF Mapper system (Bio-Rad). Electrophoresis was performed at 4.5 V cm⁻¹ and 14 °C for 48 h, with a pulse time of 120 s. Gels were stained with 1 mg mL⁻¹ ethidium bromide for 30 min before imaging. Gels were depurinated in 0.25 M HCl for 30 min, incubated in denaturation buffer (1.5 M NaCl and 0.5 M NaOH) for 1 h, and subsequently incubated in neutralisation buffer (1.5 M NaCl and 0.5 M Tris-HCl, pH 7.0) for 1 h. DNA was transferred onto Hybond-N+ membranes (Amersham) by capillary transfer in 10 x SSPE [1.5 M NaCl and 0.1 M sodium phosphate buffer (NaH₂PO₄/Na₂HPO₄), pH 7.5] for 48 h. Probes were prepared, and membranes were hybridised using the AlkPhos Direct Labeling and Detection System (Amersham), according to the manufacturer’s instructions. Chemiluminescence was detected by exposure to Hyperfilm ECL (Amersham) following incubation with CDP-Star reagent (Amersham).

### DNA probes for Southern blotting

For detection of *DHFR-TS*, a fragment encompassing the full-length gene (1,500 bp) was amplified by PCR using the forward primer 5′-ATGCCCGGGCATATGTCCAGGGCAGCTGCGAGGTTT-3′ and the reverse primer 5′-ATGCCCGGGCATATGCTATACGGCCATCTCCATCTT-3′. Telomeric sequences were detected using a 300 bp fragment containing *Trypanosoma brucei* telomeric repeats (TTAGGG), obtained by HindIII digestion of the pTEL construct (59). Before labelling, both the PCR product and the restriction fragment were purified using the illustra GFX PCR DNA and Gel Band Purification Kit (Cytiva), according to the manufacturer’s instructions.

### Quantification of DHFR-TS and PTR1 copy number by quantitative real-time PCR (qPCR)

Approximately 2.5 x 10⁷ parasites were suspended in culture medium and heated at 95 °C for 5 min. A 4-μL aliquot of the resulting lysate, equivalent to 2 x 10⁶ cells, was used as the template in a 20-μL reaction containing 1 x GoTaq qPCR Master Mix (Promega), 0.2 μL of CXR reference dye and 300 nM of each primer. Amplification and fluorescence acquisition were performed using a 7500 Real-Time PCR System (Applied Biosystems).

Cycling conditions comprised 50 °C for 2 min and 95 °C for 2 min, followed by 40 cycles of 95 °C for 15 s, 60 °C for 15 s and 72 °C for 1 min. For melting-curve analysis, amplification products were heated to 95 °C for 15 s and cooled to 60 °C for 1 min.

*DHFR-TS* was amplified using the forward primer 5′-CATCGAGACAGCGTATTGCG-3′ and the reverse primer 5′-GATGCGGGTCAGGTACACTT-3′. *PTR1* was amplified using the forward primer 5′-AGCCCTGTGTCGGAGATAGA-3′ and the reverse primer 5′-ATGCGCGAACGCCTTAATC-3′. For both targets, *G6PDH* was used as the reference gene for normalisation and amplified using the forward primer 5′-ACCGCATTGACCACTACCTC-3′ and the reverse primer 5′-GATGTTGTTCGAGTTCCAC-3′.

### DNA content analysis

Cells were harvested and washed in 1 x PBS supplemented with 5 mM EDTA. Next, cells were fixed in 70% (v/v) methanol diluted in 1 x PBS at -20 °C for 2 h. Then, cells were washed with 1 x PBS supplemented with 5 mM EDTA and incubated for 30 min at room temperature with 10 μg mL^−1^ Propidium Iodide and 10 μg mL^−1^ RNase A diluted in 1 x PBS containing 5 mM EDTA. Cells were then passed through a 35 μm nylon mesh and examined using a BD FACSCanto II flow cytometer (Becton Dickinson). Further data analysis and data processing was performed with FlowJo software.

### Detection of S-phase cells under denaturing conditions

As previously described (48), exponentially growing cells were incubated with 100 μM IdU for 30 min. After washing with 1x PBS, cells were fixed in 70% (v/v) ethanol diluted in 1 x PBS at -20 °C for 16 h. Then, cells were washed with blocking buffer [1% (w/v) BSA in 1 x PBS] and treated with 2 M HCl for 20 min at room temperature, followed by neutralisation in 0.2 M phosphate buffer (Na₂HPO₄/KH₂PO₄, pH 7.4) for 30 min at room temperature. To detect IdU incorporated during S-phase, cells were incubated for 1 h at room temperature in staining buffer [1 x PBS containing 1% (w/v) BSA and 0.2% (v/v) Tween-20] supplemented with mouse anti-BrdU antibody (clone B44; BD Biosciences; 1:300). Following washing with blocking buffer, cells were incubated for 1 h at room temperature in staining buffer containing Alexa Fluor 488-conjugated anti-mouse secondary antibody (Thermo Fisher Scientific; 1:500). Cells were washed again with blocking buffer and resuspended in 1 x PBS containing 10 μg mL⁻¹ propidium iodide and 20 μg mL⁻¹ RNase, then incubated for 30 min at room temperature.

Finally, cell suspensions were passed through a 60-μm nylon mesh (Merck Millipore) and analysed using a BD FACSCanto II flow cytometer (Becton Dickinson). Further data analysis and data processing was performed with FlowJo software.

### Detection of single-stranded DNA under non-denaturing conditions

This assay was performed as previously described (31), with minor modifications. Briefly, exponentially growing cells were incubated with 100 μM IdU for 15 h at 26 °C, transferred to fresh medium containing 10 mM HU for 15 h, and subsequently transferred to fresh, drug-free medium.

After HU removal, samples were collected at the time points indicated in the main text, washed with 1 x PBS and fixed in 70% (v/v) ethanol in 1 x PBS at −20 °C for 16 h. To detect IdU exposed within single-stranded DNA regions, cells were immunostained as described above for S-phase analysis, omitting the HCl treatment to maintain non-denaturing conditions. Further data analysis and data processing was performed with FlowJo software.

### Whole-genome sequencing, ChIP-seq and MFA-seq datasets

Whole-genome sequencing and MFA-seq datasets for *WT*, *RAD9*^+/−^ and *HUS1*^+/−^ cell lines were obtained from a previous study (30). ChIP-seq (45) and MFA-seq data were standardised by z-score transformation to facilitate comparison of relative signal patterns across experimental conditions and factors.

### Copy number variation analysis

Genome-wide CNVs were determined with bamCompare (DeepTools), using the RPKM normalization method. Pair-ended extension was employed, PCR duplicates were ignored, and regions were centered with respect to the fragment length. Fold-change was expressed as log2. Chromosome copy number was estimated from WGS read depth over annotated single-copy genes. Only properly mapped reads with mapping quality ≥30 and base quality ≥20 were considered; unmapped, secondary, supplementary, duplicate and quality-control-failed reads were excluded, and overlapping mates were counted once. Median coverage was calculated for each single-copy gene and then across all selected genes on each chromosome. Chromosome somy was estimated by normalising chromosome-level median coverage to the genome-wide median chromosome coverage, assuming that most chromosomes were disomic: somy = 2 x (median coverage / genome-wide median chromosome coverage). Continuous somy estimates were retained to capture mosaic aneuploidy.

### Genome-wide detection of candidate intrachromosomal rearrangements

Paired-end whole-genome sequencing alignments were analysed using custom scripts adapted from the Circle_finder strategy (40). Alignments were grouped by read name using SAMtools, and duplicates were marked using SAMBLASTER. Following exclusion of unmapped and secondary alignments, read pairs represented by exactly three alignment records were selected. Supplementary alignment tags were used to identify two putative split-read alignments and one contiguous mate alignment. Candidates were retained when both split-read segments mapped to the same chromosome and strand, while the contiguous mate mapped to the opposite strand of the same chromosome, with its alignment start lying strictly between the starts of the two split-read segments. Candidate intervals were defined by these split-alignment start positions and grouped by chromosome, coordinates and strand. Support was quantified as the number of distinct read-pair identifiers per interval, and separate call sets requiring at least one or two supporting read pairs were generated.

## Acknowledgements

This work was funded by Conselho Nacional de Desenvolvimento Científico e Tecnológico (CNPq) for R.O. [132367/2013-0]; Fundação de Amparo a Pesquisa do Estado de São Paulo (FAPESP) for L.R.O.T. [14/06824-9; 14/06824-8; 16/50193-8; 16/50050-2; 17/07092-9; 18/14398-0; 18/14432-3; 23/03015-0], J.A.B. [20/01883-7; 24/08412-0; 25/15819-2], S. V. [16/18191-5] and J.D.D. [13/00570-1; 14/00751-9; 25/10952-6]. Work in R.M.’s lab was supported by the Wellcome Trust [083485; 104111; 224501/Z/21/Z] and the BBSRC [BB/K006495/1; BB/M028909/1; BB/N016165/1; BB/R017166/1]. And the European Union’s Horizon 2020 research and innovation programme funded J.D.D. under the Marie Sklodowska-Curie grant agreement No 750259 [Individual Fellowship, RECREPEMLE].

## References

1. D. Dedukh, A. Krasikova, Delete and survive: strategies of programmed genetic material elimination in eukaryotes. Biol Rev Camb Philos Soc 97, 195–216 (2022).

2. G. S. Darmasaputra, L. M. van Rijnberk, M. Galli, Functional consequences of somatic polyploidy in development. Development 151 (2024).

3. P. Vande Zande, X. Zhou, A. Selmecki, The Dynamic Fungal Genome: Polyploidy, Aneuploidy and Copy Number Variation in Response to Stress. Annu Rev Microbiol 77, 341–361 (2023).

4. S. M. Christie, C. Fijen, E. Rothenberg, V(D)J Recombination: Recent Insights in Formation of the Recombinase Complex and Recruitment of DNA Repair Machinery. Front Cell Dev Biol 10, 886718 (2022).

5. I. Rzeszutek, X. X. Maurer-Alcala, M. Nowacki, Programmed genome rearrangements in ciliates. Cell Mol Life Sci 77, 4615–4629 (2020).

6. A. Tubbs, A. Nussenzweig, Endogenous DNA Damage as a Source of Genomic Instability in Cancer. Cell 168, 644–656 (2017).

7. M. R. Cosenza, B. Rodriguez-Martin, J. O. Korbel, Structural Variation in Cancer: Role, Prevalence, and Mechanisms. Annu Rev Genomics Hum Genet 23, 123–152 (2022).

8. N. K. Chunduri, Z. Storchova, The diverse consequences of aneuploidy. Nat Cell Biol 21, 54–62 (2019).

9. C. M. Brennan et al., Protein aggregation mediates stoichiometry of protein complexes in aneuploid cells. Genes Dev 33, 1031–1047 (2019).

10. J. C. Saldivar, D. Cortez, K. A. Cimprich, The essential kinase ATR: ensuring faithful duplication of a challenging genome. Nat Rev Mol Cell Biol 18, 622–636 (2017).

11. S. Saxena, L. Zou, Hallmarks of DNA replication stress. Mol Cell 82, 2298–2314 (2022).

12. S. Minocherhomji et al., Replication stress activates DNA repair synthesis in mitosis. Nature 528, 286–290 (2015).

13. N. Shaikh et al., Replication stress generates distinctive landscapes of DNA copy number alterations and chromosome scale losses. Genome Biol 23, 223 (2022).

14. L. A. Yates, X. Zhang, P. M. Burgers, DNA Damage and Replication Stress Checkpoints. Annu Rev Biochem 94, 195–221 (2025).

15. M. Day, A. W. Oliver, L. H. Pearl, Structure of the human RAD17-RFC clamp loader and 9-1-1 checkpoint clamp bound to a dsDNA-ssDNA junction. Nucleic Acids Res 50, 8279–8289 (2022).

16. S. Burza, S. L. Croft, M. Boelaert, Leishmaniasis. Lancet 392, 951–970 (2018).

17. M. Pareyn et al., Leishmaniasis. Nat Rev Dis Primers 11, 81 (2025).

18. J. M. Ubeda et al., Genome-wide stochastic adaptive DNA amplification at direct and inverted DNA repeats in the parasite Leishmania. PLoS Biol 12, e1001868 (2014).

19. Y. Sterkers, L. Lachaud, L. Crobu, P. Bastien, M. Pages, FISH analysis reveals aneuploidy and continual generation of chromosomal mosaicism in Leishmania major. Cell Microbiol 13, 274–283 (2011).

20. L. Lachaud et al., Constitutive mosaic aneuploidy is a unique genetic feature widespread in the Leishmania genus. Microbes Infect 16, 61–66 (2014).

21. G. H. Negreira et al., High throughput single-cell genome sequencing gives insights into the generation and evolution of mosaic aneuploidy in Leishmania donovani. Nucleic Acids Res 50, 293–305 (2022).

22. A. C. Ivens et al., The genome of the kinetoplastid parasite, Leishmania major. Science 309, 436–442 (2005).

23. S. A. Iantorno et al., Gene Expression in Leishmania Is Regulated Predominantly by Gene Dosage. mBio 8 (2017).

24. B. Cuypers et al., Four layer multi-omics reveals molecular responses to aneuploidy in Leishmania. PLoS Pathog 18, e1010848 (2022).

25. J. B. T. Carnielli et al., A Leishmania infantum genetic marker associated with miltefosine treatment failure for visceral leishmaniasis. EBioMedicine 36, 83–91 (2018).

26. C. A. Grace et al., Parasite Genotype Is a Major Predictor of Mortality from Visceral Leishmaniasis. mBio 13, e0206822 (2022).

27. J. D. Damasceno et al., R-loops acted on by RNase H1 influence DNA replication timing and genome stability in Leishmania. Nat Commun 16, 1470 (2025).

28. G. L. A. da Silva et al., ATR, a DNA damage kinase, modulates DNA replication timing in Leishmania major. PLoS Genet 21, e1011899 (2025).

29. J. D. Damasceno et al., Functional compartmentalization of Rad9 and Hus1 reveals diverse assembly of the 9-1-1 complex components during the DNA damage response in Leishmania. Mol Microbiol 101, 1054–1068 (2016).

30. J. D. Damasceno et al., Genome duplication in Leishmania major relies on persistent subtelomeric DNA replication. Elife 9 (2020).

31. J. D. Damasceno et al., Conditional genome engineering reveals canonical and divergent roles for the Hus1 component of the 9-1-1 complex in the maintenance of the plastic genome of Leishmania. Nucleic Acids Res 46, 11835–11846 (2018).

32. C. A. Marques, N. J. Dickens, D. Paape, S. J. Campbell, R. McCulloch, Genome-wide mapping reveals single-origin chromosome replication in Leishmania, a eukaryotic microbe. Genome Biol 16, 230 (2015).

33. J. A. Coderre, S. M. Beverley, R. T. Schimke, D. V. Santi, Overproduction of a bifunctional thymidylate synthetase-dihydrofolate reductase and DNA amplification in methotrexate-resistant Leishmania tropica. Proc Natl Acad Sci U S A 80, 2132–2136 (1983).

34. A. R. Bello, B. Nare, D. Freedman, L. Hardy, S. M. Beverley, PTR1: a reductase mediating salvage of oxidized pteridines and methotrexate resistance in the protozoan parasite Leishmania major. Proc Natl Acad Sci U S A 91, 11442–11446 (1994).

35. A. Olmo, R. Arrebola, V. Bernier, D. Gonzalez-Pacanowska, L. M. Ruiz-Perez, Co-existence of circular and multiple linear amplicons in methotrexate-resistant Leishmania. Nucleic Acids Res 23, 2856–2864 (1995).

36. J. Watson, Patient care. Aust Fam Physician 7, suppl 3-4 (1978).

37. X. Feng et al., Amplification of an alternate transporter gene suppresses the avirulent phenotype of glucose transporter null mutants in Leishmania mexicana. Mol Microbiol 71, 369–381 (2009).

38. A. Mukherjee, L. D. Langston, M. Ouellette, Intrachromosomal tandem duplication and repeat expansion during attempts to inactivate the subtelomeric essential gene GSH1 in Leishmania. Nucleic Acids Res 39, 7499–7511 (2011).

39. T. Downing et al., Whole genome sequencing of multiple Leishmania donovani clinical isolates provides insights into population structure and mechanisms of drug resistance. Genome Res 21, 2143–2156 (2011).

40. P. Kumar et al., ATAC-seq identifies thousands of extrachromosomal circular DNA in cancer and cell lines. Sci Adv 6, eaba2489 (2020).

41. D. M. Iovannisci, S. M. Beverley, Structural alterations of chromosome 2 in Leishmania major as evidence for diploidy, including spontaneous amplification of the mini-exon array. Mol Biochem Parasitol 34, 177–188 (1989).

42. O. Fernandes, V. K. Murthy, U. Kurath, W. M. Degrave, D. A. Campbell, Mini-exon gene variation in human pathogenic Leishmania species. Mol Biochem Parasitol 66, 261–271 (1994).

43. R. Inga et al., Relation between variation in copy number of ribosomal RNA encoding genes and size of harbouring chromosomes in Leishmania of subgenus Viannia. Mol Biochem Parasitol 92, 219–228 (1998).

44. S. Martinez-Calvillo et al., Genomic organization and functional characterization of the Leishmania major Friedlin ribosomal RNA gene locus. Mol Biochem Parasitol 116, 147–157 (2001).

45. J. A. Black et al., Genome-wide mapping of replication stress response factors reveals the management of polycistronic transcription, replication initiation, and responses to replication stress in Leishmania. Nucleic Acids Res 54 (2026).

46. J. D. Damasceno, V. S. Nunes, L. R. Tosi, LmHus1 is required for the DNA damage response in Leishmania major and forms a complex with an unusual Rad9 homologue. Mol Microbiol 90, 1074–1087 (2013).

47. L. Glover, D. Horn, Trypanosomal histone gammaH2A and the DNA damage response. Mol Biochem Parasitol 183, 78–83 (2012).

48. J. D. Damasceno et al., Conditional knockout of RAD51-related genes in Leishmania major reveals a critical role for homologous recombination during genome replication. PLoS Genet 16, e1008828 (2020).

49. M. M. Genois et al., Roles of Rad51 paralogs for promoting homologous recombination in Leishmania infantum. Nucleic Acids Res 43, 2701–2715 (2015).

50. M. C. Laffitte et al., Formation of linear amplicons with inverted duplications in Leishmania requires the MRE11 nuclease. PLoS Genet 10, e1004805 (2014).

51. G. H. Ngo, L. Balakrishnan, M. Dubarry, J. L. Campbell, D. Lydall, The 9-1-1 checkpoint clamp stimulates DNA resection by Dna2-Sgs1 and Exo1. Nucleic Acids Res 42, 10516–10528 (2014).

52. F. L. Tsai, M. Kai, The checkpoint clamp protein Rad9 facilitates DNA-end resection and prevents alternative non-homologous end joining. Cell Cycle 13, 3460–3464 (2014).

53. D. Salim et al., DNA replication stress restricts ribosomal DNA copy number. PLoS Genet 13, e1007006 (2017).

54. F. Dumetz et al., Modulation of Aneuploidy in Leishmania donovani during Adaptation to Different In Vitro and In Vivo Environments and Its Impact on Gene Expression. mBio 8 (2017).

55. G. Bussotti et al., Leishmania Genome Dynamics during Environmental Adaptation Reveal Strain-Specific Differences in Gene Copy Number Variation, Karyotype Instability, and Telomeric Amplification. mBio 9 (2018).

56. R. A. Burrell et al., Replication stress links structural and numerical cancer chromosomal instability. Nature 494, 492–496 (2013).

57. T. Wilhelm et al., Mild replication stress causes chromosome mis-segregation via premature centriole disengagement. Nat Commun 10, 3585 (2019).

58. T. E. Ellenberger, J. E. Wright, A. Rosowsky, S. M. Beverley, Wild-type and drug-resistant Leishmania major hydrolyze methotrexate to N-10-methyl-4-deoxy-4-aminopteroate without accumulation of methotrexate polyglutamates. J Biol Chem 264, 15960–15966 (1989).

59. L. H. Van der Ploeg, A. Y. Liu, P. Borst, Structure of the growing telomeres of Trypanosomes. Cell 36, 459–468 (1984).

