## Supplemental Data for "Replication-stress response shapes *Leishmania* genome plasticity through non-equivalent RAD9 and HUS1 functions"

### Supplementary Figure 1

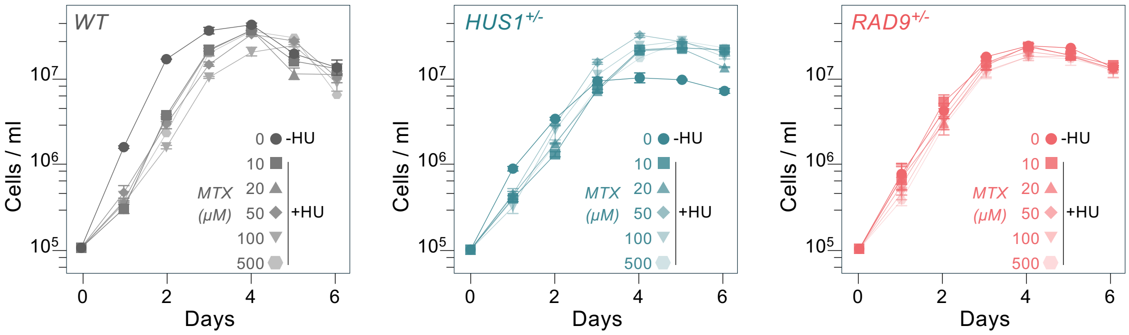


**Supplementary Figure 1.** Growth profiles of *WT*, *HUS1*$^{+/-}$ and *RAD9*$^{+/-}$ cells subjected to MTS selection after HU treatment as indicated in main Fig. 2A.

### Supplementary Figure 2

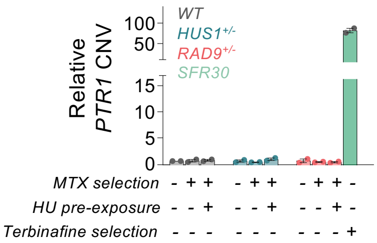


**Supplementary Figure 2.** Quantitative PCR measurement of *PTR1* copy number in MTX-selected populations with or without prior HU exposure. Ct values were normalised to G6PDH and expressed relative to Ct value from unselected WT. Bars represent means ± SEM from two independent biological replicates. *SFR30*, cell line obtained from step-wise selection with terbinafine and was used as a positive control for *PTR1* amplification.

### Supplementary Figure 3

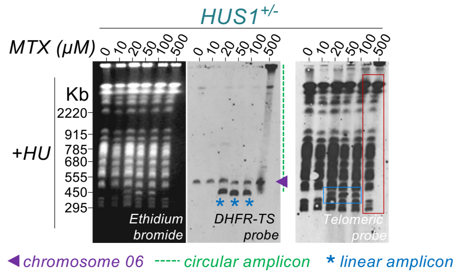


**Supplementary Figure 3.** Pulsed-field gel electrophoresis (PFGE) and Southern-blot analysis of the indicated MTX-selected populations. Intact chromosomes and extrachromosomal DNA molecules were visualised by ethidium-bromide staining prior to transferring. The membrane shown in main Fig. 1B (*HUS1*$^{+/-}$ pre-exposed to HU) was striped and re-hybridised with a telomeric probe. Purple arrowheads, signal of intact chromosome 6. Blue asterisks, linear amplicons. Green dashed lines, circular amplicons. Blue rectangle, telomere-positive amplicons. Red rectangle, absence of telomere-positive smear.

### Supplementary Figure 4

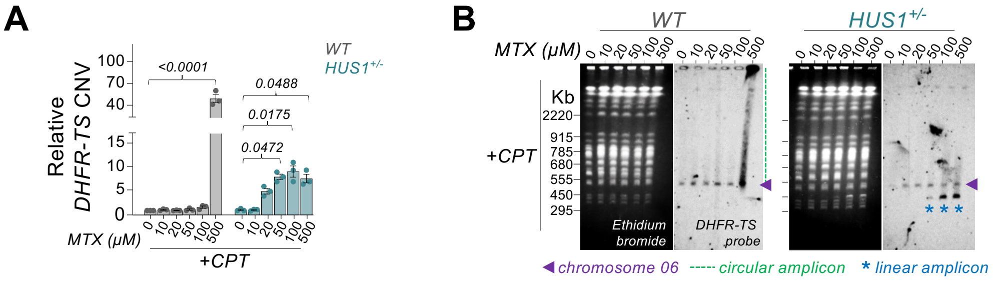


**Supplementary Figure 4. (A)** Quantitative PCR measurement of *DHFR-TS* copy number in MTX-selected populations with prior CPT exposure. Ct values were normalised to G6PDH and expressed relative to Ct value from unselected WT. Bars represent means ± SEM from three independent biological replicates. *P* values were determined with Friedman multiple comparisons test. **(B)** Pulsed-field gel electrophoresis (PFGE) and Southern-blot analysis of the indicated MTX-selected populations with prior CPT exposure. Intact chromosomes and extrachromosomal DNA molecules were visualised by ethidium-bromide staining prior to transferring. Membranes were hybridised with a *DHFR-TS* probe. Purple arrowheads, signal of intact chromosome 6. Blue asterisks, linear amplicons. Green dashed lines, circular amplicons.

### Supplementary Figure 5

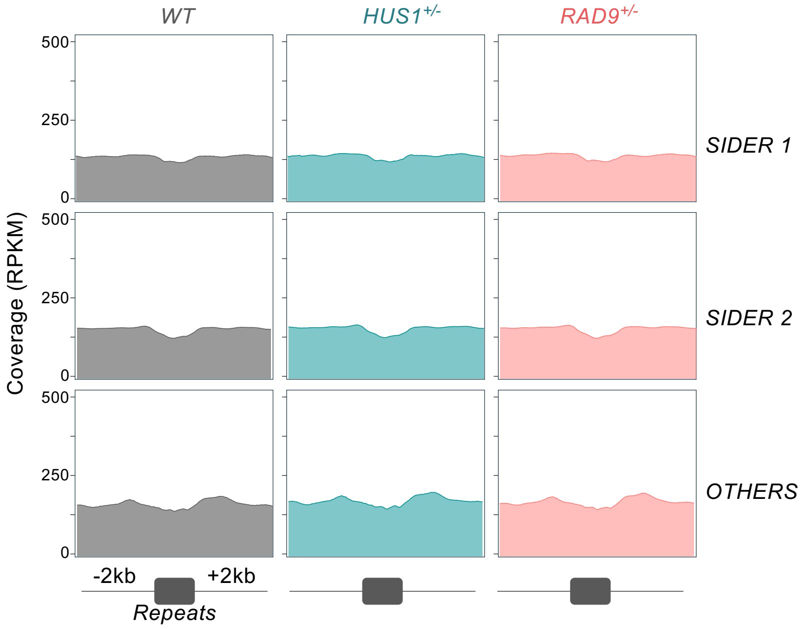


**Supplementary Figure 5.** Read depth across repetitive DNA sequences. Repeats were grouped according to the classification by *Ubeda, J. M. et al. PLoS biology, 2014*.
